# enclone: precision clonotyping and analysis of immune receptors

**DOI:** 10.1101/2022.04.21.489084

**Authors:** David B. Jaffe, Payam Shahi, Bruce A. Adams, Ashley M. Chrisman, Peter M. Finnegan, Nandhini Raman, Ariel E. Royall, FuNien Tsai, Thomas Vollbrecht, Daniel S. Reyes, Wyatt J. McDonnell

## Abstract

Half a billion years of evolutionary battle forged the vertebrate adaptive immune system, an astonishingly versatile factory for molecules that can adapt to arbitrary attacks. The history of an individual encounter is chronicled within a clonotype: the descendants of a single fully rearranged adaptive immune cell. For B cells, reading this immune history for an individual remains a fundamental challenge of modern immunology. Identification of such clonotypes is a magnificently challenging problem for three reasons:

- <u>The cell history is inferred rather than directly observed</u>: the only available data are the sequences of V(D)J molecules occurring in a sample of cells.
- <u>Each immune receptor is a pair of V(D)J molecules</u>. Identifying these pairs at scale is a technological challenge and cannot be done with perfect accuracy—real samples are mixtures of cells and fragments thereof.
- <u>These molecules can be intensely mutated</u> during the optimization of the response to particular antigens, blurring distinctions between kindred molecules.

It is thus impossible to determine clonotypes exactly. All solutions to this problem make a trade-off between sensitivity and specificity; useful solutions must address actual artifacts found in real data.

We present **enclone**^1^, a system for computing approximate clonotypes from single cell data, and demonstrate its use and value with the 10x Genomics Immune Profiling Solution. To test it, we generate data for 1.6 million individual B cells, from four humans, including deliberately enriched memory cells, to tax the algorithm and provide a resource for the community. We analytically determine the specificity of **enclone**’s clonotyping algorithm, showing that on this dataset the probability of co-clonotyping two unrelated B cells is around 10^-9^. We prove that using only heavy chains increases the error rate by two orders of magnitude.

**enclone** comprises a comprehensive toolkit for the analysis and display of immune receptor data. It is ultra-fast, easy to install, has public source code, comes with public data, and is documented at bit.ly/enclone. It has three “flavors” of use: (1) as a command-line tool run from a terminal window, that yields visual output; (2) as a command-line tool that yields parseable output that can be fed to other programs; and (3) as a graphical version (GUI).

## Introduction

T and B cells are part of the adaptive immune system of vertebrates, playing an essential role in long-term interactions with infection, cancer, symbiosis, and autoimmunity. For this, their ability to target specific antigens is enabled by exquisitely specialized heterodimeric proteins: T cell receptors (TCRs) and B cell receptors (BCRs). TCRs are membrane-bound, whereas BCRs exist in both membrane-bound and secreted forms, known as antibodies.

T and B cells access a vast range (estimated to lie between 10^8^ and 10^16^) of possible receptors by editing their own genomes, first rearranging one chain (TRB or heavy/IGH) in a V(D)J recombination process, and then rearranging the other chain (TRA or light) in a VJ recombination process, yielding a “fully rearranged cell” [reviewed in: Jung 2006]. In B cells, receptor sequence editing continues through somatic hypermutation. The complete set of descendants of a fully rearranged cell comprise a **clonotype^2^**, and thus the entirety of an individual’s T or B cells are partitioned into clonotypes. Clonotypes are sometimes also called **lineages**.

A holy grail of immunology is the ability to read out and decipher an individual’s clonotypes. This has deep implications both for basic science and medicine, including the development of vaccines, antibody therapeutics, and cell therapies. The starting point for identifying clonotypes is a laboratory process for determining V(D)J sequences. The most straightforward approach is bulk sequencing [Barennes 2021, Bashford-Rogers 2014]. This samples the TRB or IGH sequences of an individual, and separately, the TRA or IGK/IGL sequences. This method is fundamentally limited as it cannot directly connect the two, though it is dramatically less expensive than other approaches and remains a valuable tool. The sole implementation of this method which can pair the two, PAIR-seq [Howie 2015], does not directly observe chain pairs and uses a formulation of the birthday problem to predict pairing with a considerable false positive rate. Conversely, single cell-based methods (see <u>References</u>) retain this chain-chain pairing information and thus in principle have power to reconstruct the true architecture of a clonotype.

Our goal here is the accurate and complete determination of clonotypes to the extent that is possible using a limited sample of the immune repertoire. Thus, we start with single cell data. We next posit the following minimal representation of a clonotype: the full nucleotide sequence, extending from the start codon at the beginning of the V to the end of the J for each chain of a cell. **We denote this as V..J.** Note that this is exactly the minimum information required to synthesize an antibody from BCR data for functional testing. Real T and B cells can have two TRA/light chains or, rarely, two TRB/heavy chains due to allelic inclusion, and this information should be retained and used by the clonotyping algorithm.

There are three fundamental reasons why this problem is challenging and why, at least for B cells, the solution can be only an approximation. The first is that the only available data are observed sequences of V(D)J molecules from a sample of cells, which is assumed to be a random and miniscule sampling of the true repertoire. The second is that physical samples nominally consisting of single cells are in fact mixtures of single cells, containing fragments and molecules thereof, down to individual mRNA molecules. A droplet attempting to encapsulate a single cell will inevitably include other material, and its characteristics will vary depending on the particular sample. For example, fragmentation of a *single* plasma cell, bloated with V(D)J mRNA, can lead to widespread contamination (**Figure 16**). The third is that somatic hypermutation in B cells clouds the delineation between clonotypes.

**Figure 1.**
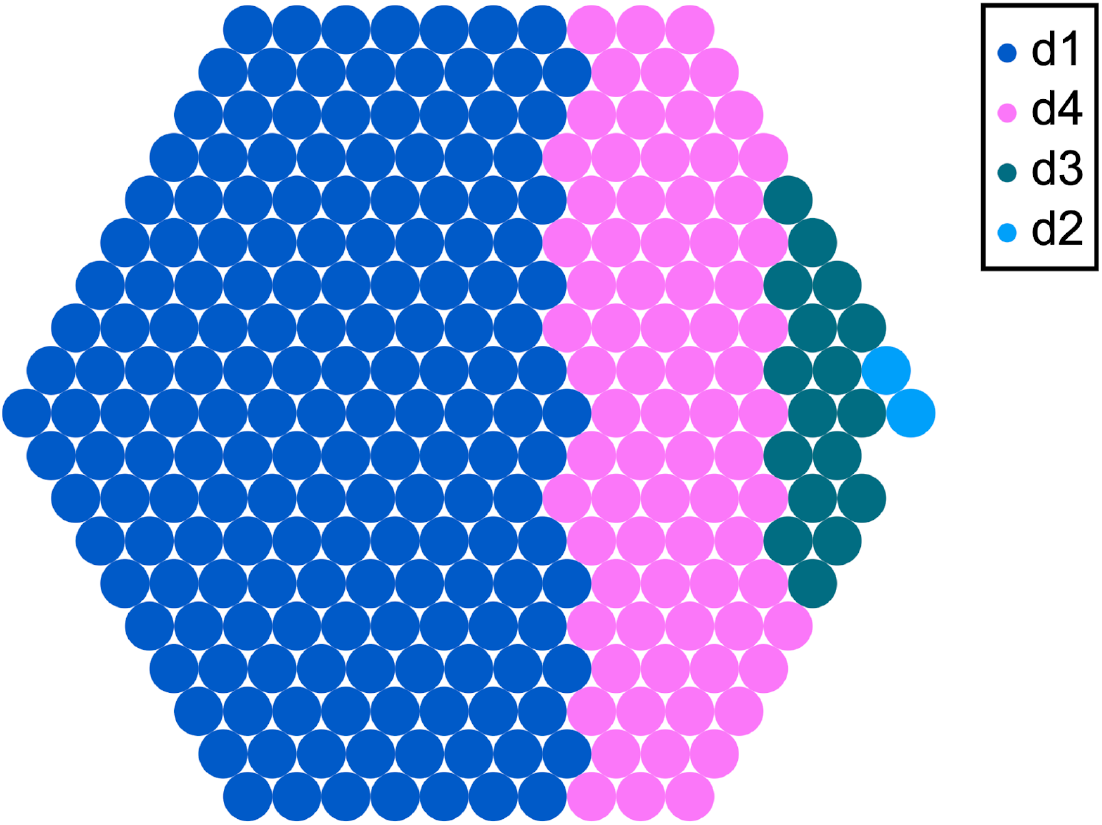
The error rate can be concentrated in a few clonotypes. The reported error rate is based on counting cell pairs. The response to “single error events” is thus quadratic and can dominate. For example, when we run **enclone** using the non-default setting `MAX_LOG_SCORE=10`, more errors are visible. These errors are concentrated in a few mixed clonotypes. Here we show the worst case, which is a clonotype using *IGHV3-9* and including CDRH3=CIKDILPGGADSW. This error contributes 18,201 to the total error score of 29,500 (62%). The above plot was generated using

~~~
enclone BCR=@test MIN_CHAINS_EXACT=2 MIX_DONORS LVARSP=donors,dref BUILT_IN
MAX_LOG_SCORE=10 CDR3=CIKDILPGGADSW
HONEY=out=fp.example1.svg,color=catvar,donors_cell,maxcat:
~~~ See subsequent sections for explanation of this command.

**Figure 2.**
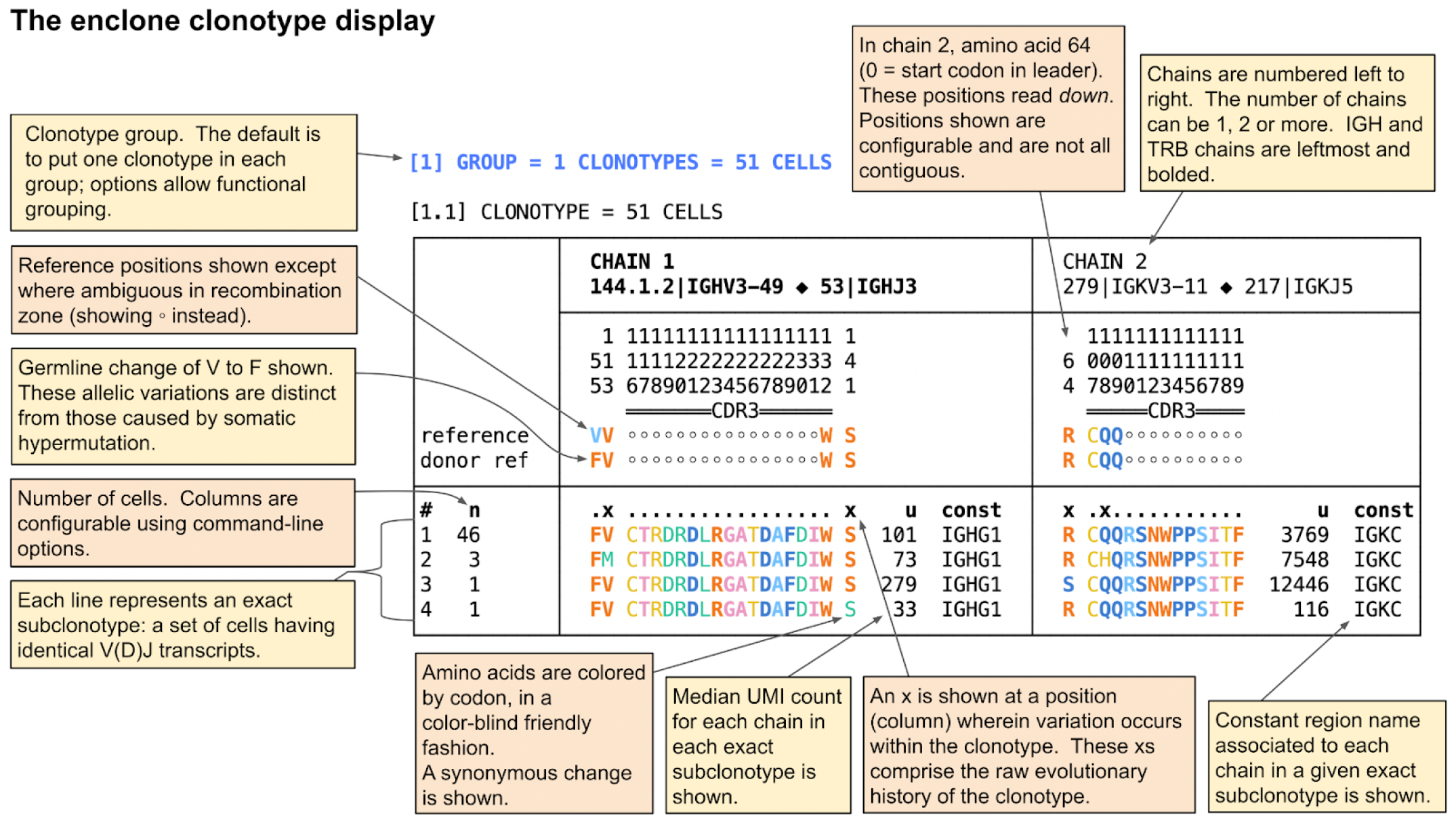
The default **enclone** display of a clonotype, which can be modified in many ways using command-line arguments. This clonotype has four exact subclonotypes, comprising a total of 51 cells.

**Figure 3.**
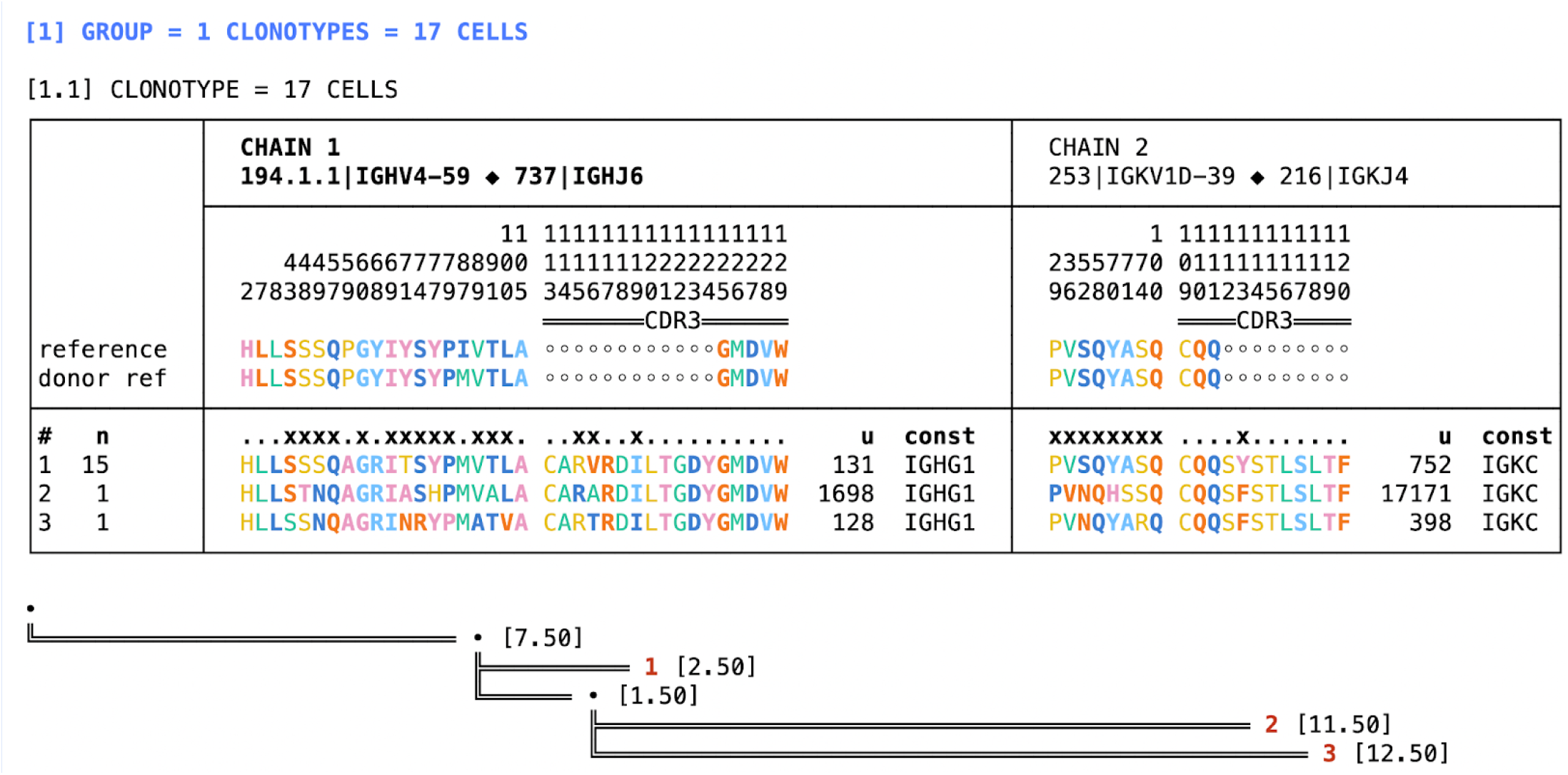
Phylogenetic tree. The output of the command

~~~
enclone BCR=123085 CDR3=CARVRDILTGDYGMDVW TREE
~~~ is exhibited. In the tree, red numbers next to each leaf refer to exact subclonotypes, and bracketed numbers represent distances (measured in nucleotides). Much more detail about this display and the underlying algorithms can be found [here](https://10xgenomics.github.io/enclone/pages/auto/tree.html) on the **enclone** site.

**Figure 4.**
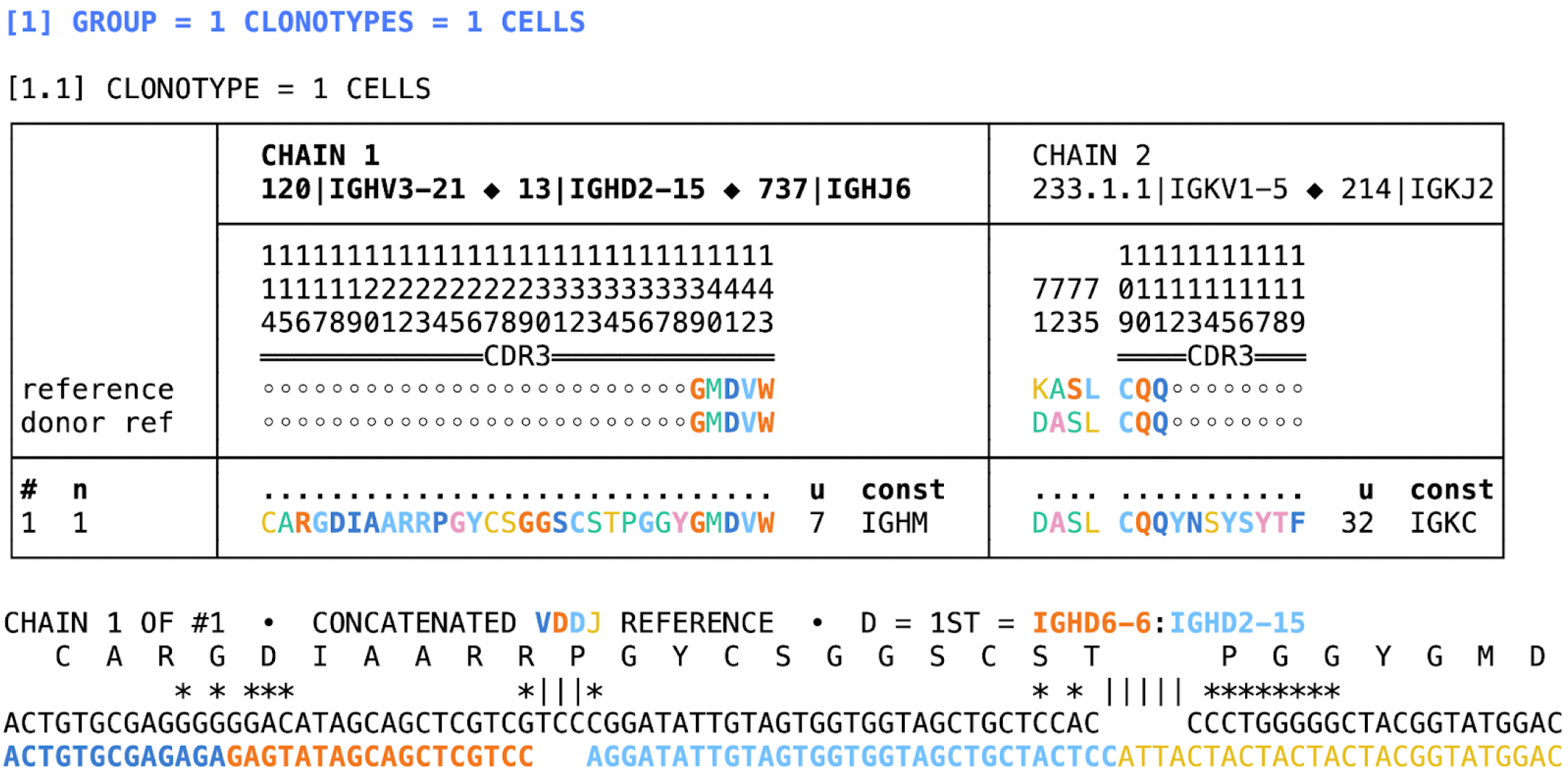
Junction region display and VDDJ. The output of the command

~~~
enclone BCR=85333 CDR3=CARGDIAARRPGYCSGGSCSTPGGYGMDVW JALIGN1
~~~ is shown. At the bottom of the display, two rows of nucleotides are shown:

- The top row is part of the heavy chain contig (representing the expressed sequence).
- The bottom row exhibits part of the concatenation of four reference sequences VDDJ. One observes in the two rows a match of the contig junction region to the reference sequences for both **IGHD6-6** and **IGHD2-15**.

**Figure 5.**
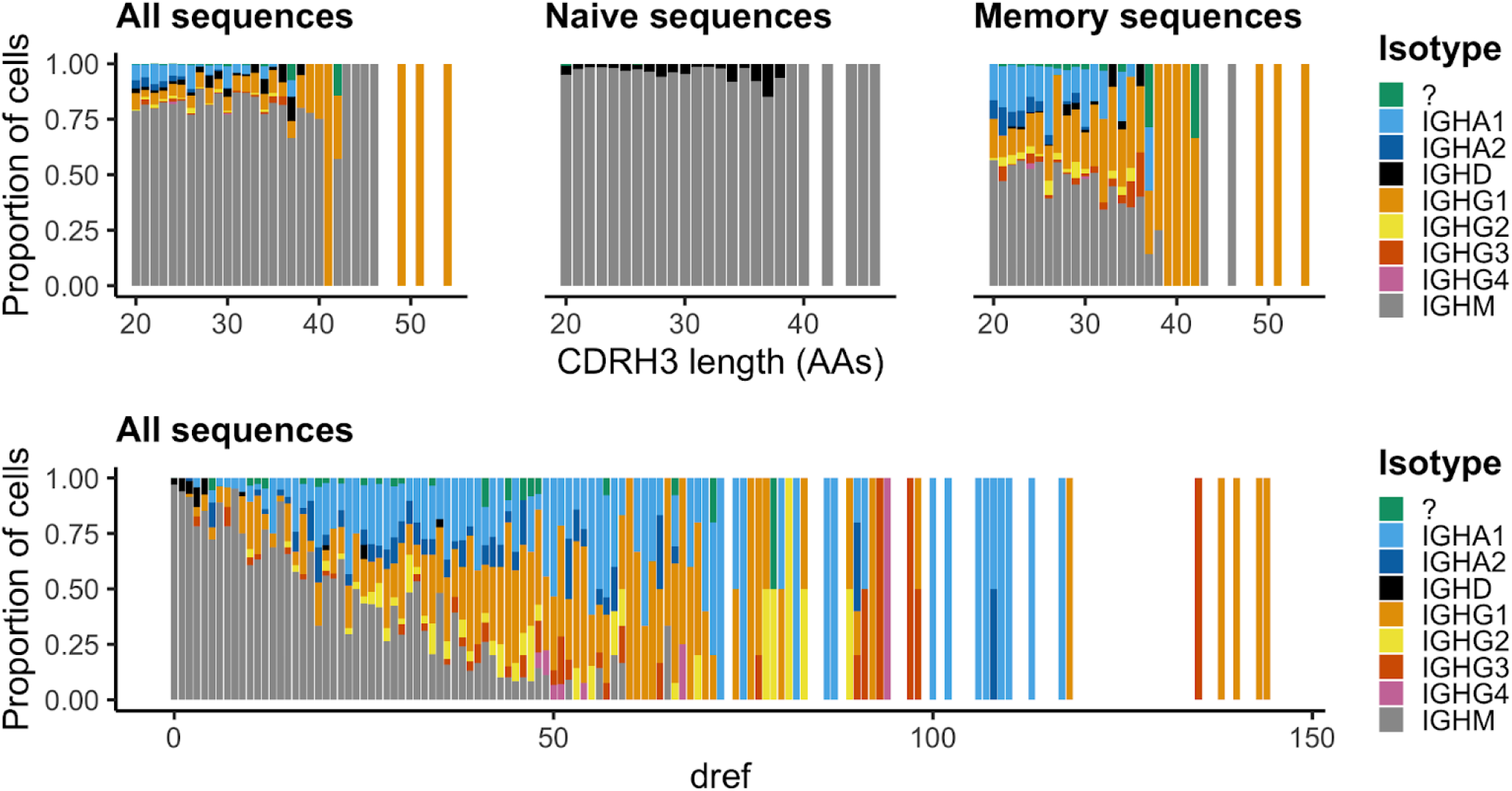
Class switching in VDDJ antibodies. <u>Top left panel</u>: the proportion of cells (Y axis) with a given CDRH3 length (X axis) is shown for all VDDJ clonotypes in the test dataset. <u>Top middle panel</u>: same for all VDDJ clonotypes with dref = 0. <u>Top right panel</u>: same for all VDDJ clonotypes with dref > 0. <u>Bottom panel</u>: the proportion of cells (Y axis) with a given dref (X axis) is shown for all VDDJ clonotypes in the test dataset. <u>Reproduction</u>: The following command generates the source data for this visualization:

~~~
enclone BCR=@test BUILT_IN D_SECOND NOPRINT PCELL MIN_CHAINS_EXACT=2
CHAINS_EXACT=2 PCOLS=cdr3_len1,cdr3_aa1,const1,dref POUT=my_output.csv
~~~

**Figure 6.**
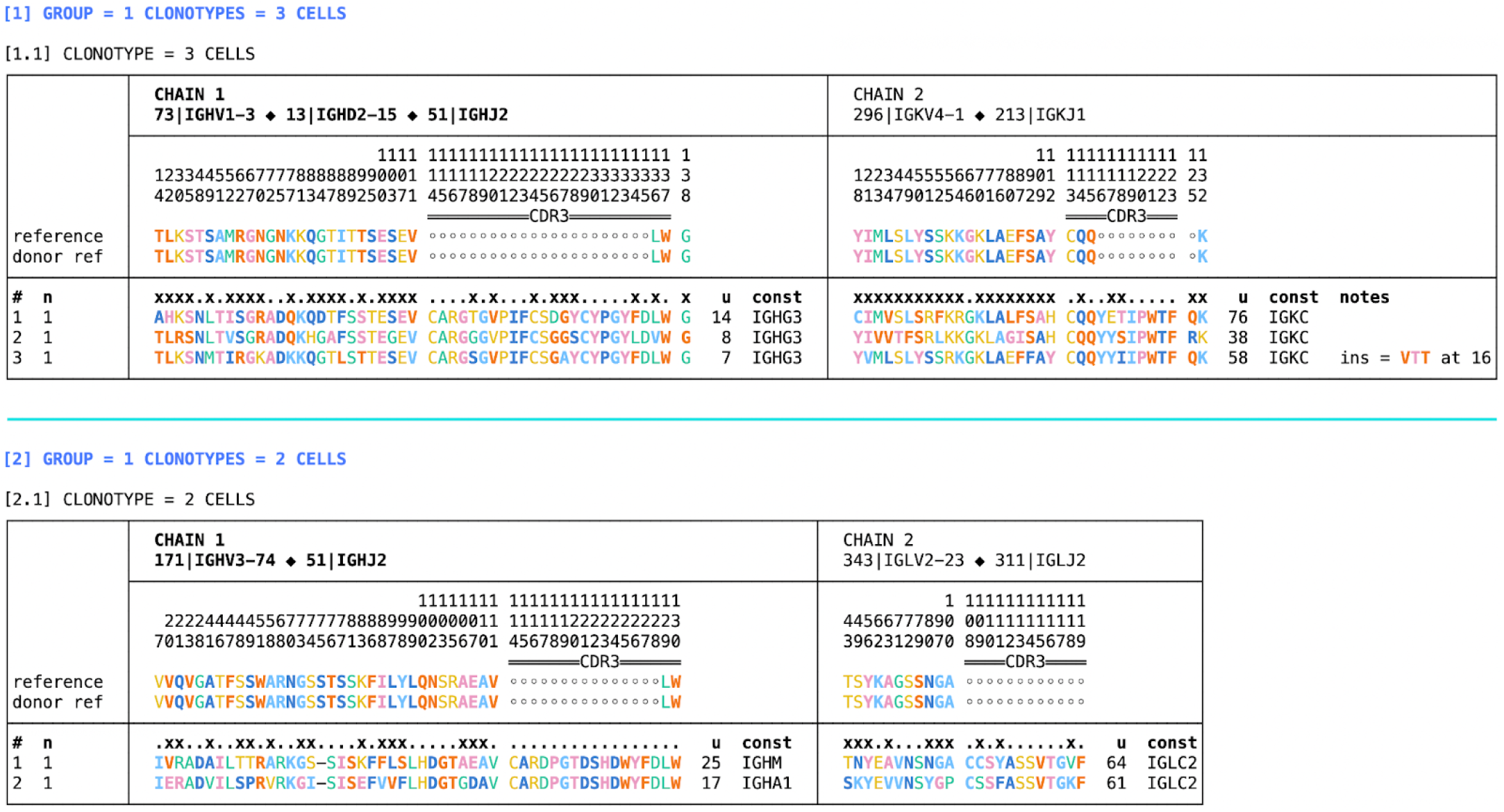
SHM insertions and deletions in clonotypes. This command

~~~
enclone BCR=40945,40961,46032,47215,47216 LVARS=n
CDR3=“CARDPGTDSHDWYFDLW|CARGSGVPIFCSGAYCYPGYFDLW”
~~~ yields the above output. In the first clonotype, there are three cells, the last of which has a three amino acid insertion in its light chain. In the second clonotype there are two cells, both of which have a single amino acid deletion in the heavy chain, denoted by `-`.

**Figure 7.**
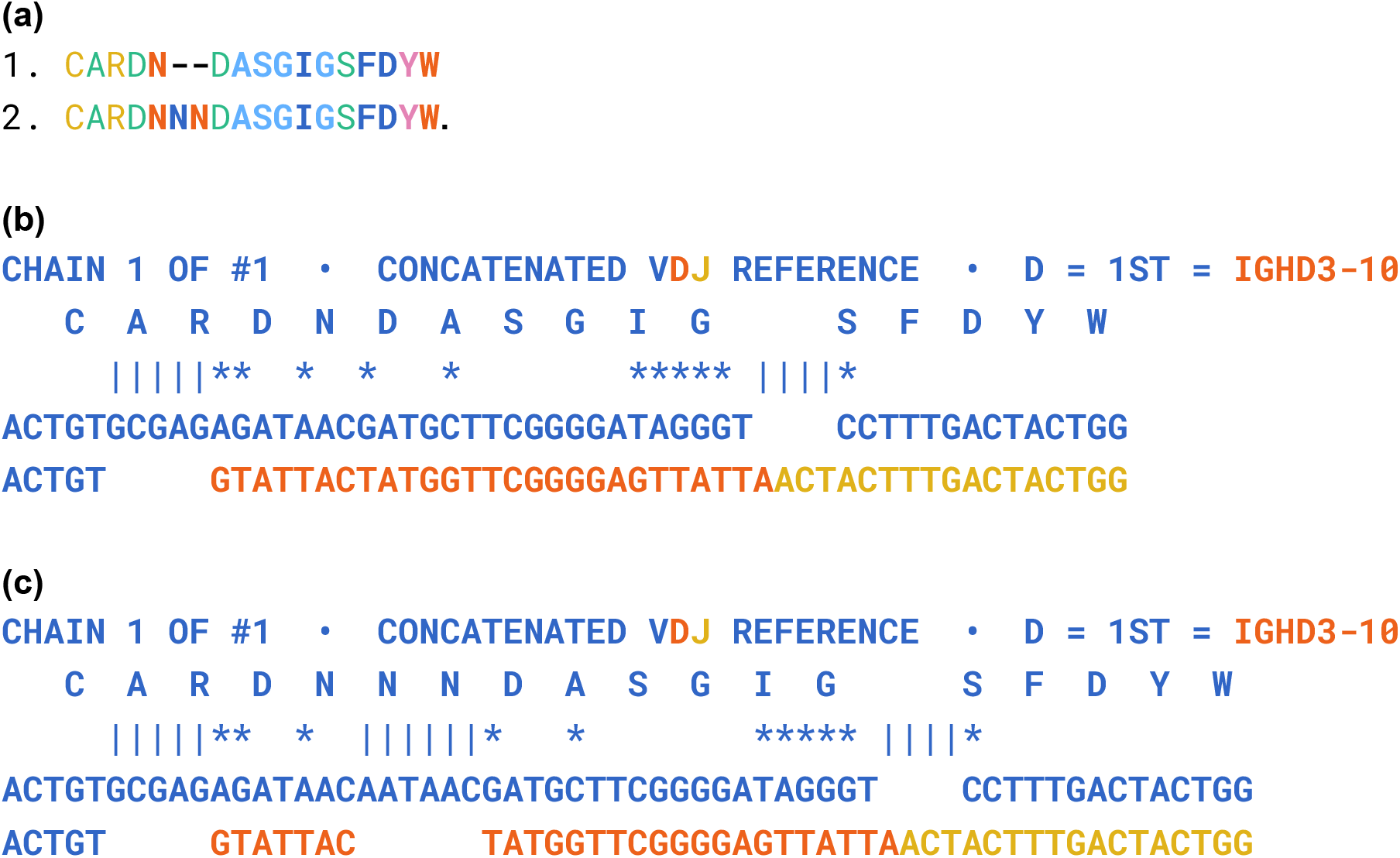
Possible SHM insertion in CDRH3. **(a)** These two CDRH3 sequences are observed in the test data, for the same donor. **(b)** Automatically generated alignment for sequence 1 (using the option JALIGN1). **(c)** Manual reconstruction of alignment for sequence 2, including putative SHM insertion. The two alignments are identical except for the insertion, and the identity of all the substitutions supports the notion that the insertion might be attributable to SHM.

**Figure 8.**
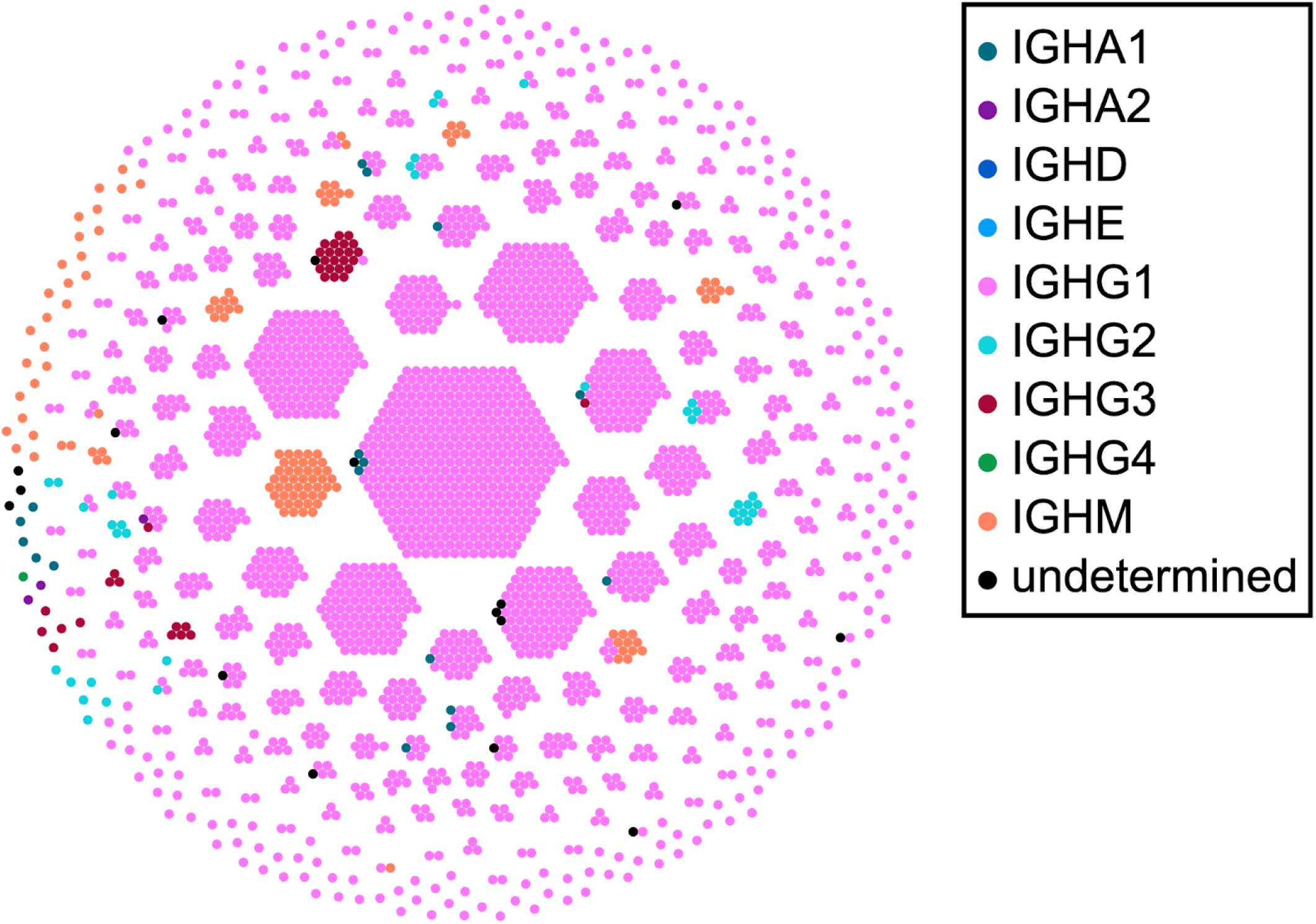
Honeycomb plot. Each dot is a cell, and each hexagonal cluster is a clonotype. We show this for the combination of two datasets from the same donor, generated using the command

~~~
enclone BCR=123085,123089 MIN_CHAINS_EXACT=2 NOPRINT PLOT_BY_ISOTYPE=plot.svg.
~~~ It does not generate output to the terminal because of the `NOPRINT` option, but does create a file `plot.svg`, as shown above. The option `MIN_CHAINS_EXACT=2` suppresses exact subclonotypes having only one chain. The latter is one of many filtering options that can be supplied to **enclone**.

**Figure 9.**
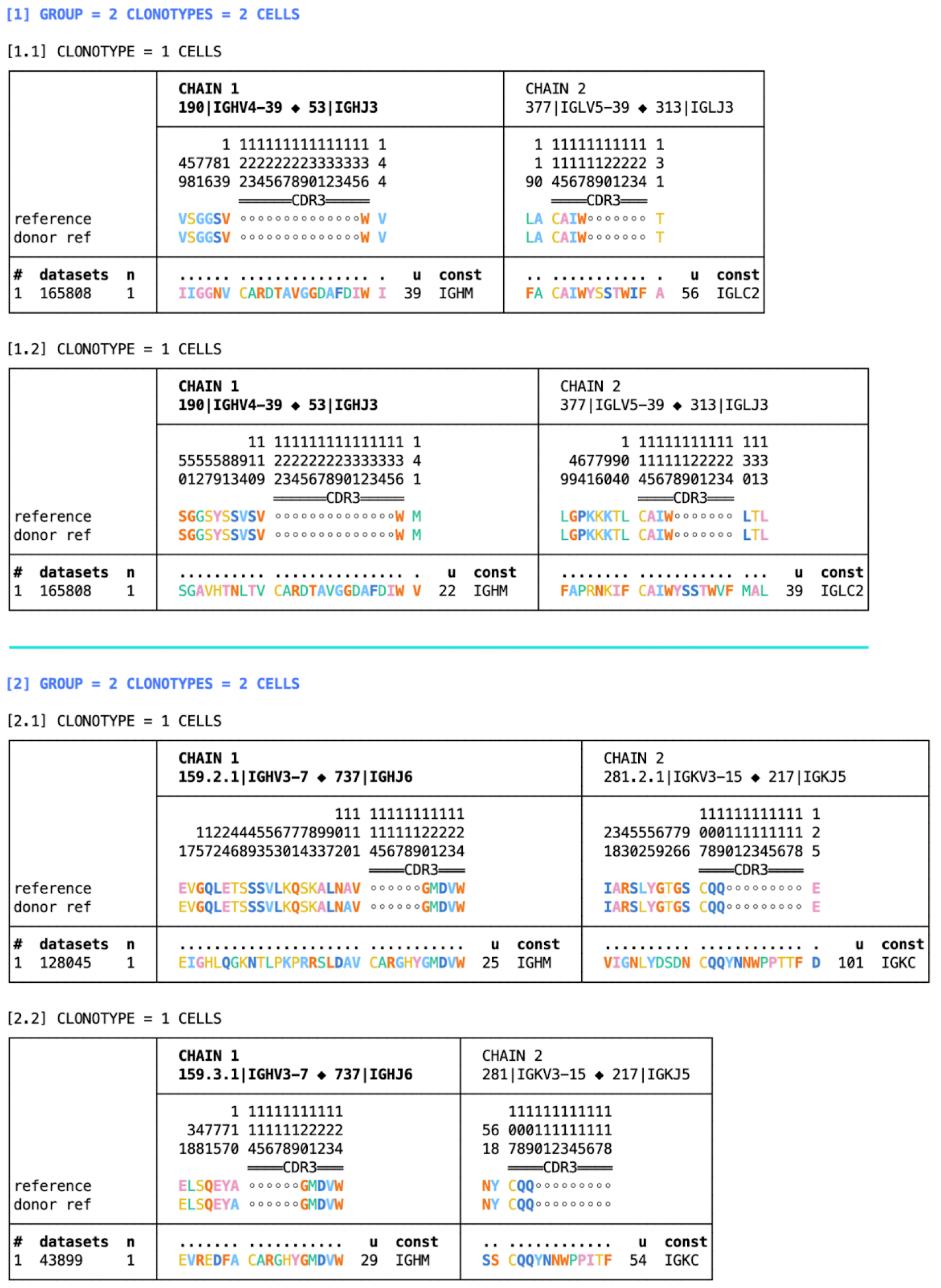
Two examples of symmetric grouping. This shows the output of

~~~
enclone GROUP=vj_refname,cdr3_aa_heavy?80% CHAINS=2 BCR=“165808;128045;43899”
CDR3=“CARGHYGMDVW|CARDTAVGGDAFDIW”.
~~~ In the first group, there are two constituent clonotypes, from the same donor. This *could* be an example of a false negative for the clonotype joining algorithm. In the second group, there are two constituent clonotypes, from different donors.

**Figure 10.**
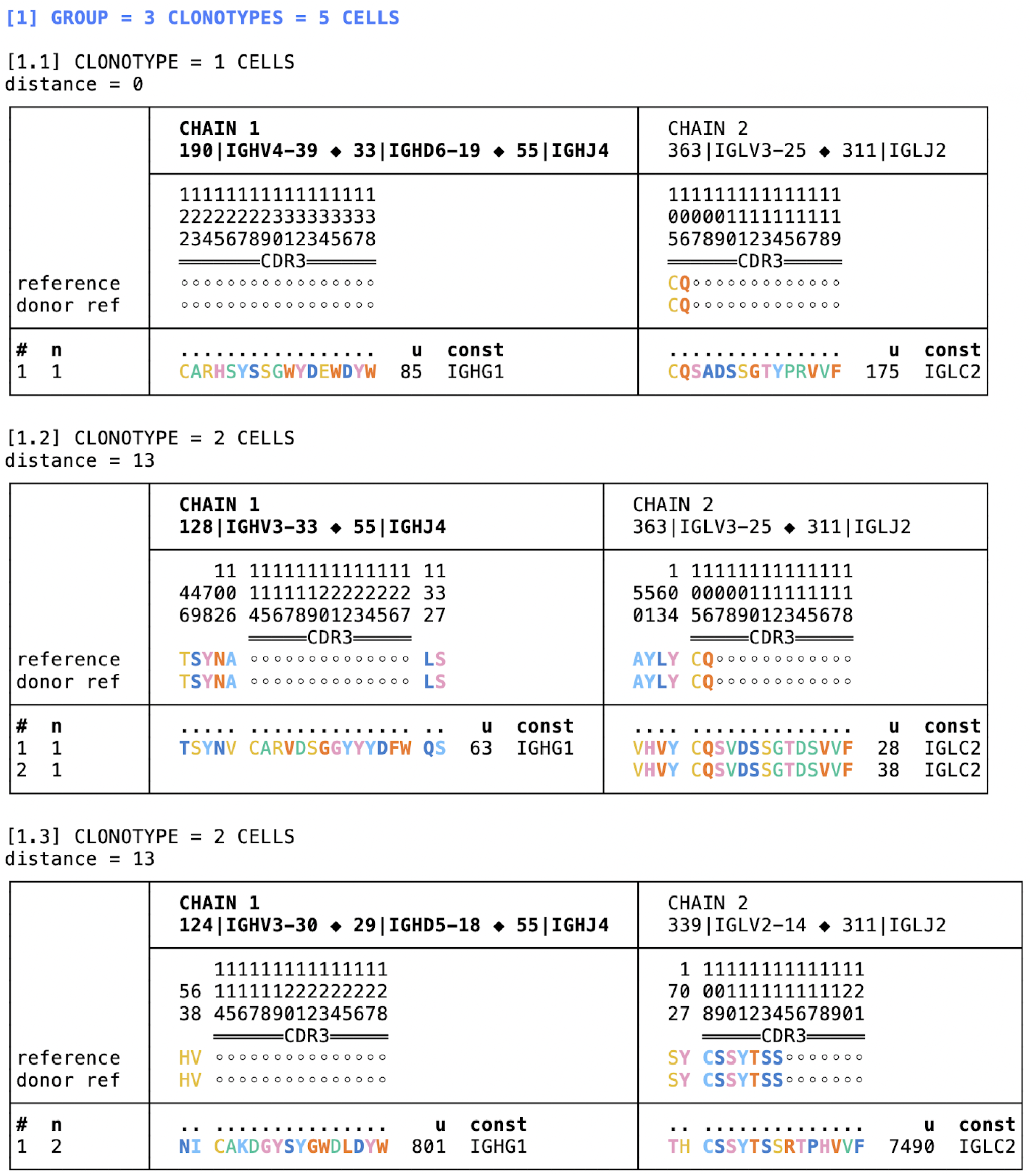
Example of asymmetric grouping. We show the output of

~~~
enclone BCR=123085 AGROUP AG_CENTER=from_filters CDR3=CARHSYSSGWYDEWDYW
AG_DIST_FORMULA=cdr3_edit_distance AG_DIST_BOUND=top=2.
~~~ For each clonotype, its two nearest neighbors by CDR3 edit distance are displayed. Using the CDR3= argument, we restrict the display to show one group.

**Figure 11.**
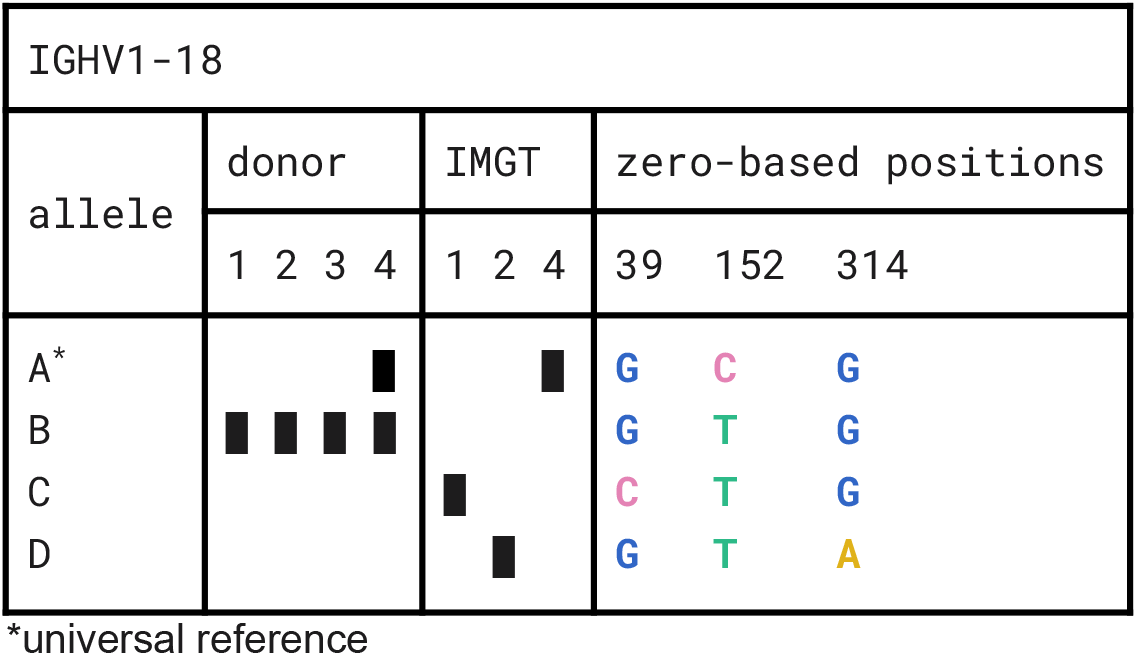
Alleles for IGHV1-18. Four alleles (A-D) are displayed. Of these, two (A-B) are found in the test data, three (A, C, D) are found in IMGT, and one (A) is found in the 10x V(D)J reference. The alleles differ at three positions. Interestingly, allele B is present in all donors, and homozygous in three, but is not in IMGT.

**Figure 12.**
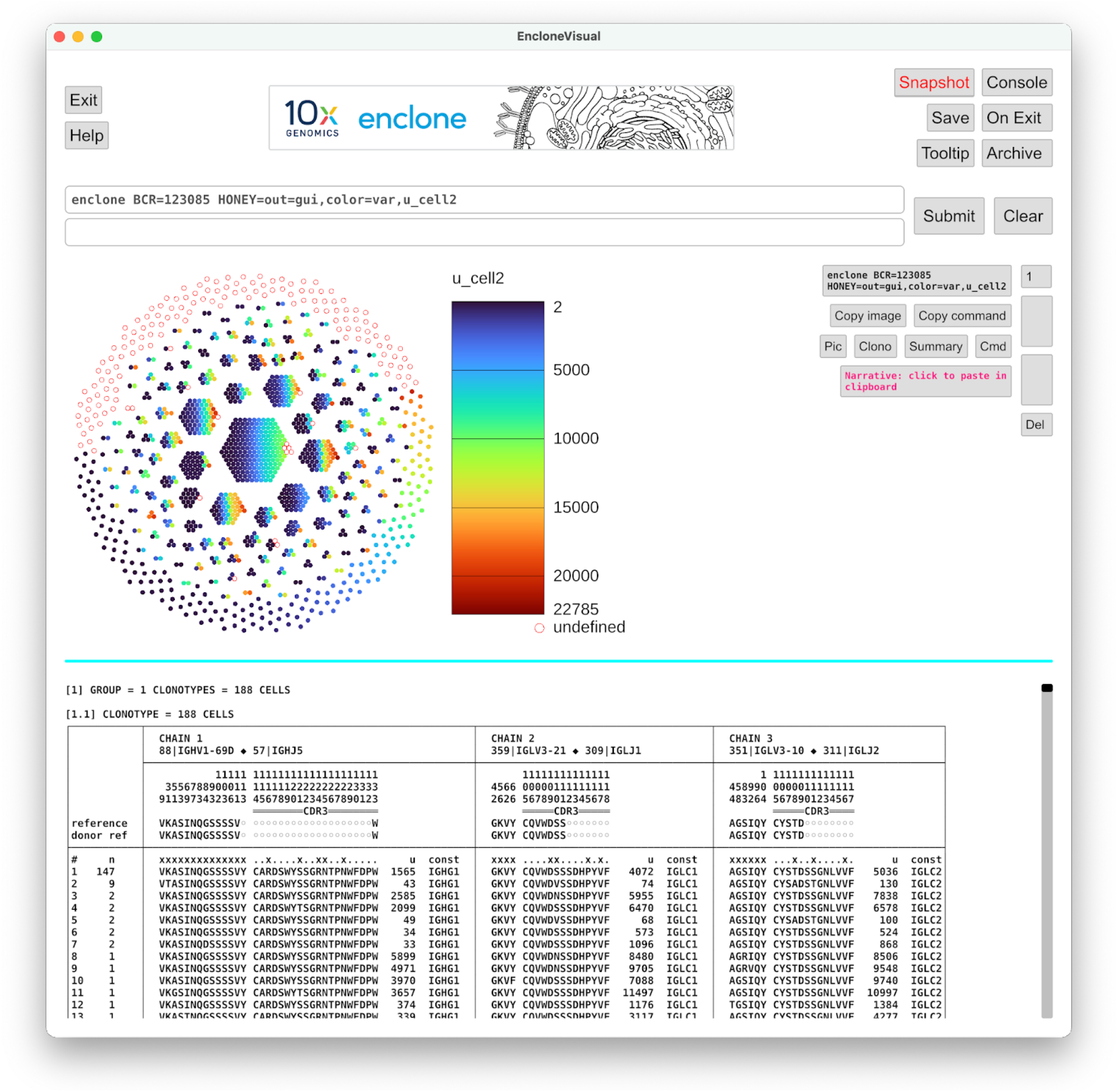
Honeycomb plot in which cells are colored by the value of a variable. Each cell is colored by the second chain UMI count. **enclone** has many other variables that might be used for this purpose.

**Figure 13.**
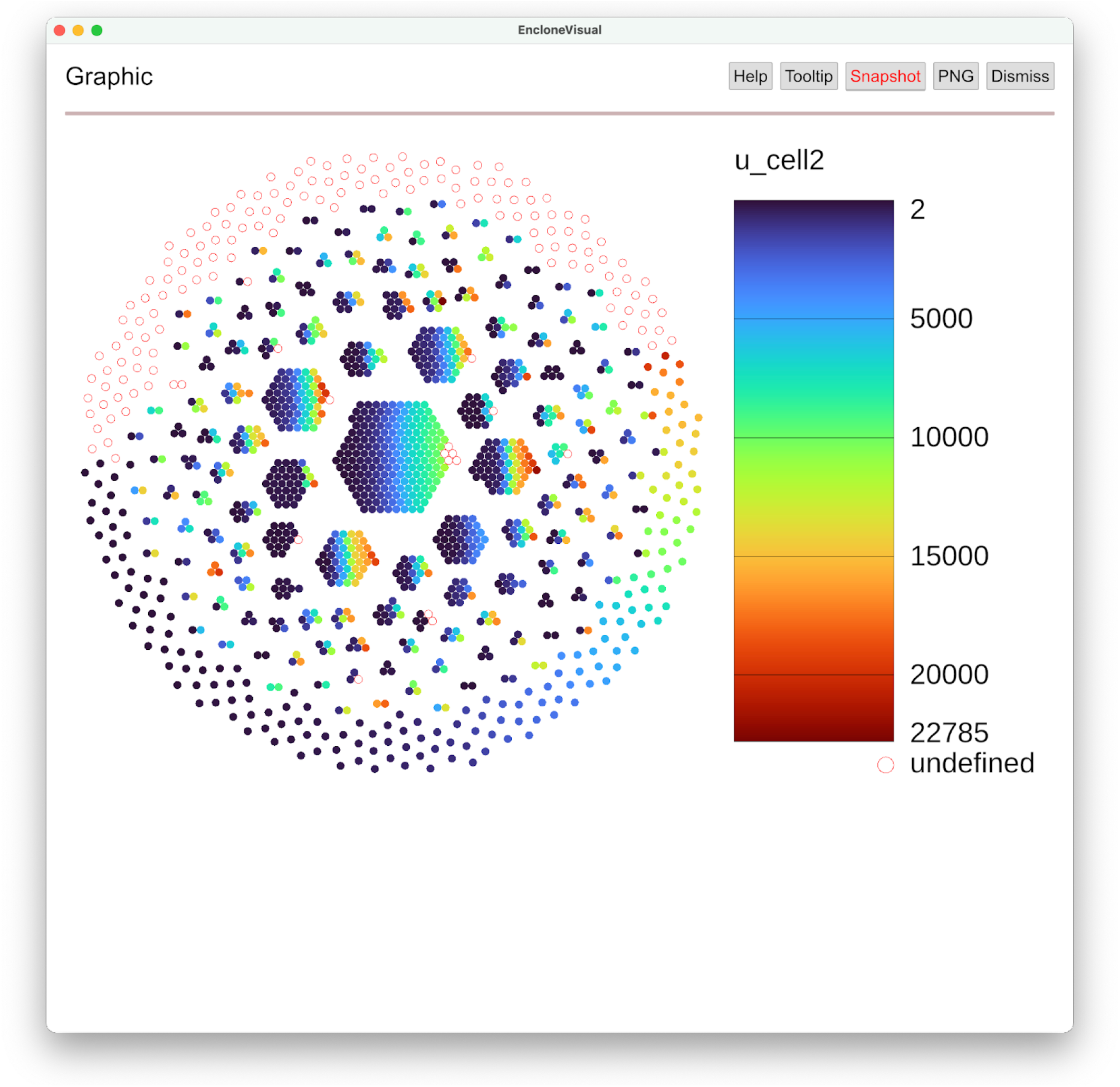
Expansion of view in enclone visual to show just the graphical object. See previous figure. Pushing the Pic button in **enclone visual** results in this display.

**Figure 14.**
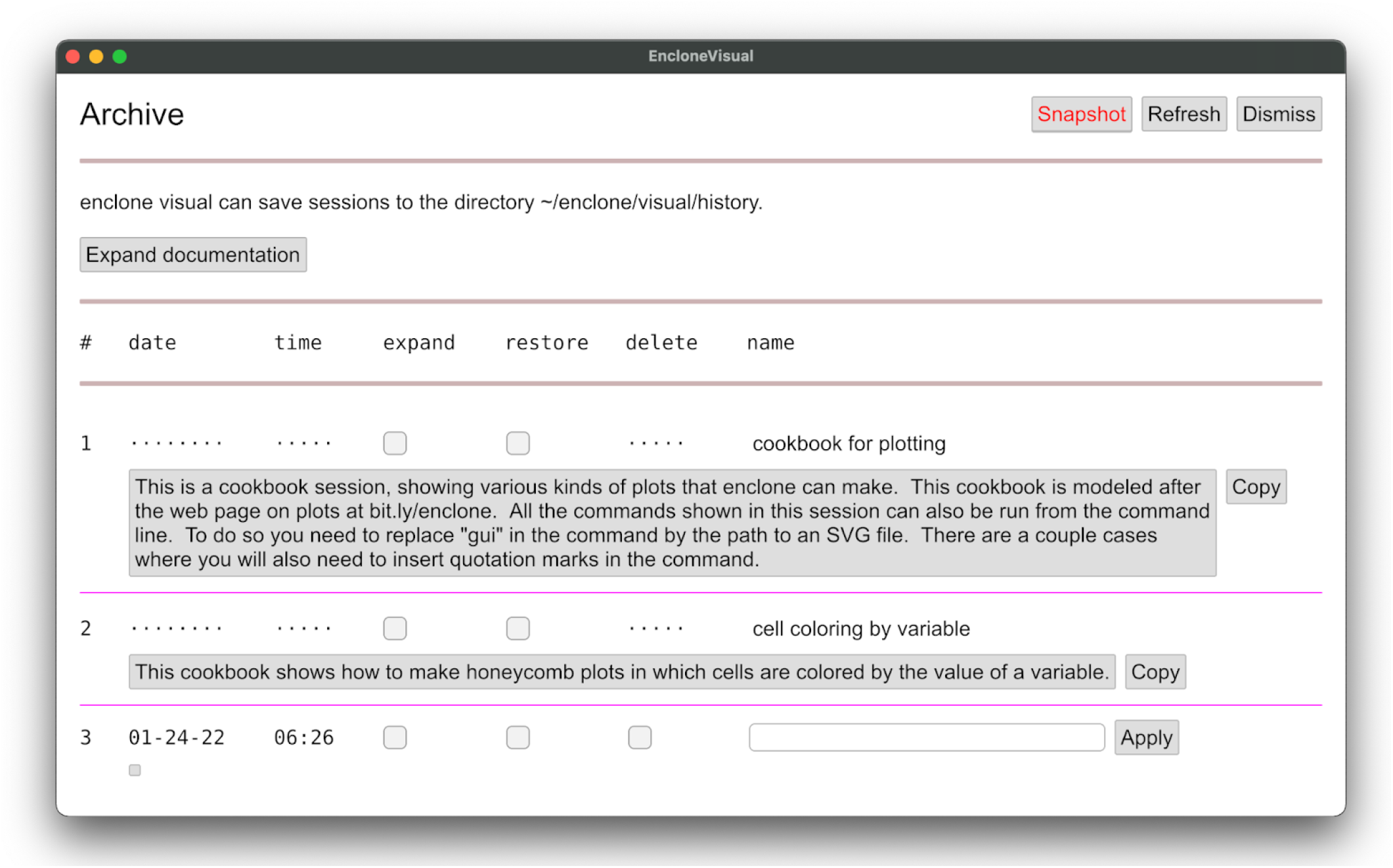
The Archive page. Previously saved sessions and built-in cookbooks are displayed.

**Figure 15.**
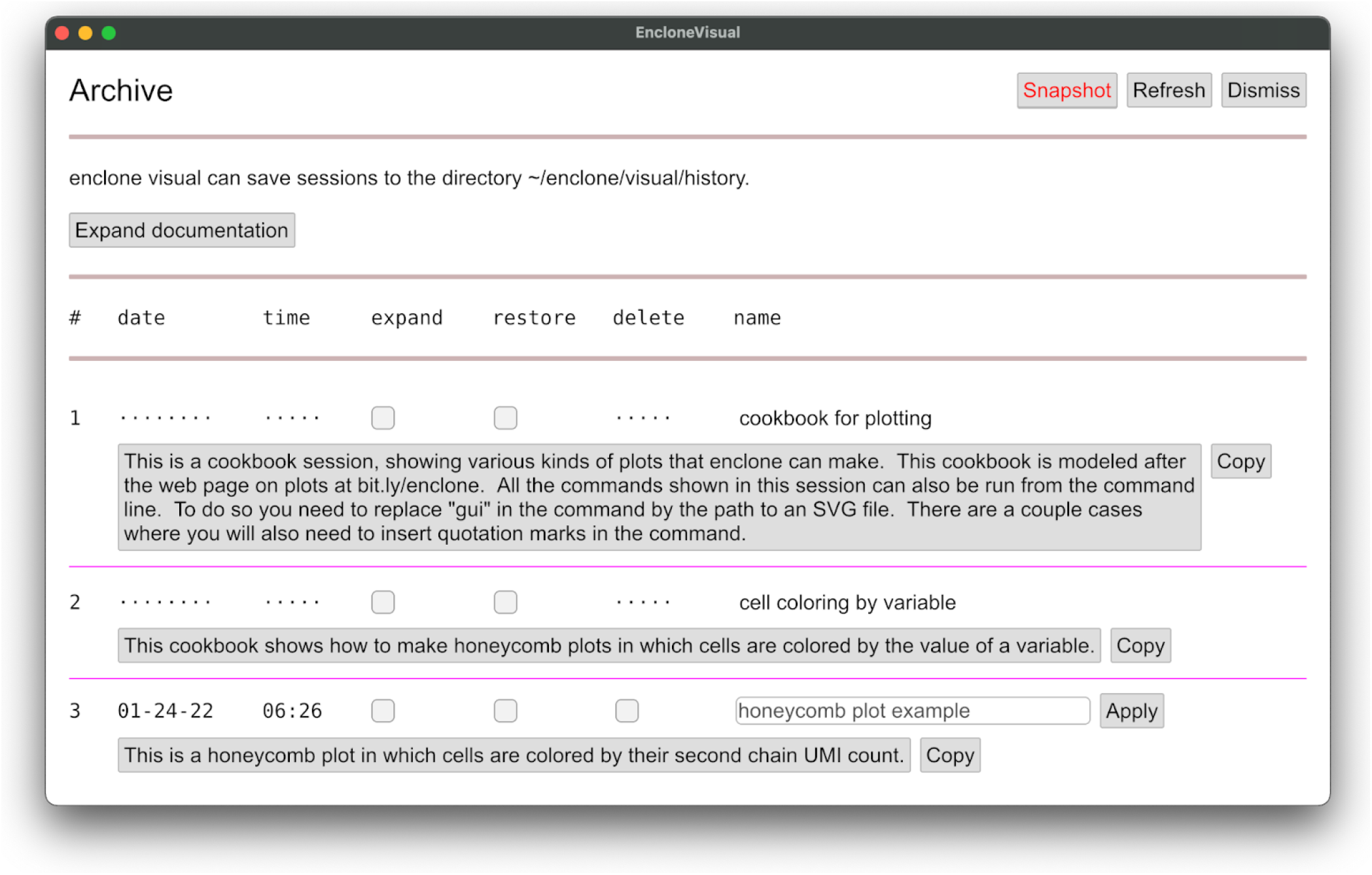
Editing the Archive page. Here we show the page after a title and narrative have been added to entry 3.

**Figure 16.**
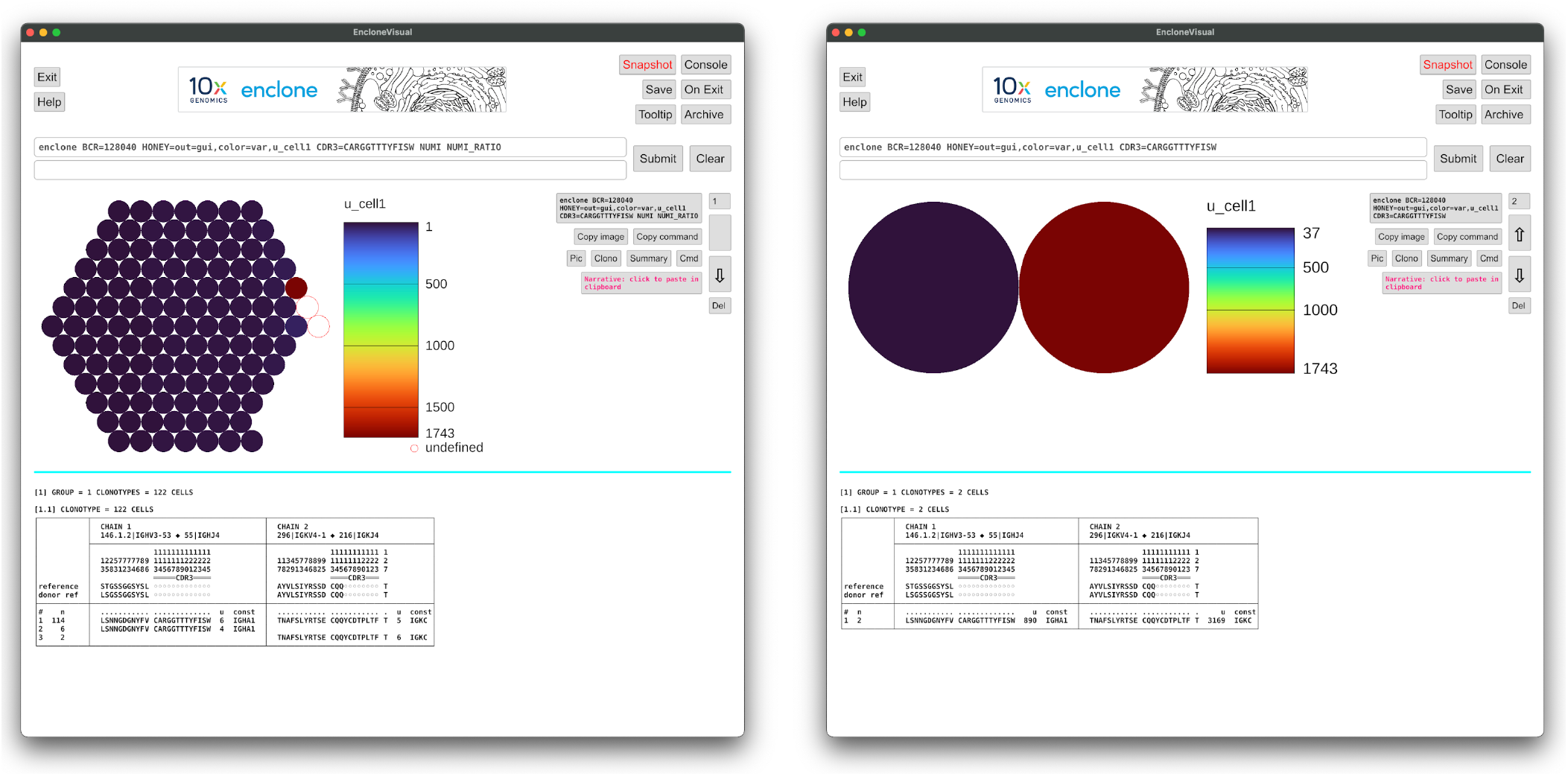
Before and after filtering. Before filtering based on UMI counts, **122** cells are reported for the clonotype having the given CDR3 (left). After filtering, only **2** cells are reported (right). Probably only **1** cell was actually present, and it was a plasma cell. The other “cells” likely represent droplets (GEMs) containing only fragments of the true “parent” cell, and have very low UMI counts as compared to the parent.

Widely used methods for B cell clonotyping are based on V/J gene concordance for either the heavy and light chains, or the heavy chain alone, together with a given nucleotide percent identity on the CDR3 regions. The <u>References</u> survey algorithms that have been used for the general problem, including recent examples of such use.

### § Assessment of clonotyping sensitivity and specificity for B cells

There is no clonotyping ground truth: examination of VDJ sequences alone cannot perfectly determine their true clonotypes. For this reason, there is no extant method for determining the sensitivity and specificity of clonotyping algorithms. We puzzled over myriad examples where we were unsure of the true answer and became confident that (lacking ground truth) we could not solve the problem by simulation. A way out could be to generate single cell data capturing both VDJ and mitochondrial mRNA [Miller 2022]. One could also engineer cell-intrinsic barcodes [Bowling 2020: Leeper 2021], but not in humans. Alternatively, since some cells are heterozygous for V or J genes, this might be exploited, however this information would be clouded by somatic hypermutation (SHM).

Finally, one can clonotype combined data from multiple individuals, and since cells from two individuals cannot lie within a single true clonotype, use this to approximate specificity [Nouri 2018 <u>Front Immunol</u>].

Our approach uses such data. We define the clonotyping error rate to be the fraction of *pairs* of cells from different individuals that are clonotyped together. This approach is “quadratically” sensitive to errors, as for example, a clonotype containing 100 cells from each of two individuals would contribute 10,000 to the numerator. In an alternative approach this might be scored as a single error (constant counting), or 100 errors (linear counting), but we think the quadratic approach yields a more meaningful metric, as large incorrect clonotypes are particularly misleading. Note that our approach is sensitive to small numbers of events.

The computed error rate depends on the datasets used and can only be used to compare algorithmic methods. For a given method one can also compute the number of pairs of cells from the *same* donor that are clonotyped together. This is a sum of true and false joins. It may be used to roughly compare methods but should not be confused with sensitivity. The computed error rate is based on cross-donor comparisons and should be correlated with intradonor clonotype mixing (true errors), but it is not the same. If it were the same, one could “subtract out inferred false joins” to yield “true sensitivity” and indeed we attempted this, but decided the approach was specious, because depending on assumptions, one can get *negative* sensitivity.

### § enclone

In this work we introduce a system for computing and working with clonotypes called **enclone**. We defer to subsequent text to describe the method, and here preview the results. We also generate and make available test data for 1.6 million B cells from four individuals, enriched for memory cells. Using the default settings, the error rate^3^ of **enclone** on these data is roughly 10^-9^.

We also compare to other clonotyping heuristics for single cell data, finding a significant advantage over the ones that we test. Importantly, we apply widely used heavy chain-only heuristics to the same data (as one might do for actual bulk data) and we find that such methods have ~100-fold higher error rate. This directly demonstrates the tremendous advantages of single cell immune repertoire sequencing methods and clonotyping methods utilizing paired sequences.

We note that one finding of this work is that sharing between donors in B cell clonotypes computed by **enclone** is rare. This appears to conflict with published results for public clonotypes, though this is explained by the use of less stringent definitions. See below, Functional grouping of B cell clonotypes.

We have described **enclone** as a clonotyping algorithm, though it is a general toolkit for studying and visualizing clonotypes^4^. It is documented in detail at bit.ly/enclone. It is very fast. It includes a command-line version (beta software) and a visual version (alpha software). The Rust source code is publicly available, as are sample datasets, including those used in this work. Improvements and contributions are most welcome. We note that **enclone** is licensed for use in connection with 10x Genomics data, though open source contributions can be made by creating separate Rust crates, and calling those from **enclone** itself (we have done this in several cases). **enclone** is our best effort, and we think it highly useful, but much more could be done with the community’s help.

The remainder of this paper consists of, first, a demonstration of clonotyping properties (**Results**), followed by a demonstration of the capabilities of **enclone**, and finally, Methods and supplementary information.

## Results

**enclone** was originally optimized based on extensive data, mostly from 10x internal protocol optimization, and also including data shared by customers, often containing edge cases not previously observed. For this work, we generated single cell V(D)J test data using six 10x Genomics Chromium X HT chip kits, comprising **1.6M B cells from 4 individuals**. The data were enriched for memory B cells to increase the clonotyping challenge. The effective size of the data was much larger than any we had previously seen. We learned new things from these data and added a <u>step</u> to the clonotyping algorithm.

We then assessed the sensitivity and specificity of several clonotyping algorithms on the totality of the data (**Table 1**). The algorithms are listed below:

1. The **enclone <u>clonotyping algorithm</u>**, and variants of it obtained by perturbing a single optional argument MAX_LOG_SCORE.
2. The partis algorithm (see <u>References</u>).
3. The algorithm **BASIC:n%.** Two cells are joined in a clonotype if they have the same heavy chain V gene assignment, the same heavy chain J gene assignment, the same light chain V gene assignment, the same light chain J gene assignment, at least n% nucleotide identity on CDRH3, and at least n% nucleotide identity on CDRL3.
4. The algorithm **BASIC_H:n%.** Two cells are joined in a clonotype if they have the same heavy chain V gene assignment, the same heavy chain J gene assignment, and at least n% nucleotide identity on CDRH3.

**Table 1.** BCR clonotyping performance of several methods.

| Heuristic | Mixed clonotypes | Error rate on interdonor comparisons | Intradonor merges (true and false) | Relative intradonor merges |
| --- | --- | --- | --- | --- |
| MLS: 5 (default) | 78 | $2.65 \times 10^{-9}$ | 2,383,558 | 1.000 |
| MLS: 10 | 259 | $33.27 \times 10^{-9}$ | 2,525,479 | 1.060 |
| MLS: 15 | 723 | $87.93 \times 10^{-9}$ | 2,668,842 | 1.120 |
| MLS: 20 | 1,800 | $207.41 \times 10^{-9}$ | 2,810,411 | 1.179 |
| partis | 450,371 | $3019.32 \times 10^{-9}$ | 2,804,940 | 1.176 |
| BASIC: 91% | 206 | $2.98 \times 10^{-9}$ | 2,256,375 | 0.947 |
| BASIC: 90% | 367 | $7.05 \times 10^{-9}$ | 2,457,566 | 1.031 |
| BASIC: 80% | 6,409 | $144.04 \times 10^{-9}$ | 3,131,915 | 1.314 |
| BASIC: 70% | 28,057 | $1981.18 \times 10^{-9}$ | 3,944,298 | 1.655 |
| BASIC_H: 92% | 5,563 | $70.24 \times 10^{-9}$ | 2,230,328 | 0.936 |
| BASIC_H: 91% | 7,986 | $146.08 \times 10^{-9}$ | 2,355,870 | 0.988 |
| BASIC_H: 90% | 11,055 | $312.89 \times 10^{-9}$ | 2,552,472 | 1.071 |
| BASIC_H: 88% | 18,048 | $1418.21 \times 10^{-9}$ | 3,175,961 | 1.332 |

**Heuristic:** the clonotyping join heuristic that was used
- MLS: MAX\_LOG\_SCORE, an enclone parameter with default 5 - partis: We ran partis in seeded “fast” paired loci mode and did not retain its contig annotations. - BASIC: instead join using V+J gene concurrence and CDR3 nucleotide similarity at given percent - BASIC\_H: same but use only heavy chain
**Mixed clonotypes:** number of clonotypes containing cells from multiple donors
**Error rate:** number of different-donor pairs of cells that were co-clonotyped, divided by interdonor comparisons
**Intradonor merges (true and false):** number of same-donor pairs of cells that were co-clonotyped
**Relative interdonor merges:** intradonor merges, normalized to 1.000 for top line

Details of how these methods were carried out are described in <u>Reproduction of results</u>. Algorithms 3 and 4 are simple and widely used (some papers are annotated as such in <u>References</u>). Algorithm 2 (partis) is a state-of-the-art method. We note that there are other sophisticated methods, but in part because of the size of our data, running any one of them was a nontrivial challenge. Thus we attempted to run the SCOPer algorithm [Nouri 2018 <u>Bioinformatics</u>; Nouri 2020], and see Methods, but terminated it when it had not finished after twelve days. Algorithms such as this may be highly performant on smaller datasets but it was not feasible for us to test them.

### § Basic observations and interpretation

#### How to read Table 1

The mixed clonotypes and error rate columns tell us about specificity. Conversely the intradonor merges column is a <u>sum of true and false merges</u>. It is not a measure of sensitivity! However, at fixed specificity, in principle, more intradonor merges should correspond to higher sensitivity.

#### Best performance

**enclone** (first line) reports the best specificity. The closest one can get (amongst algorithms tested) is BASIC: 91%, which has somewhat lower specificity and at least 5% lower sensitivity. However, note that BASIC: 80% has an observed error rate that is 50 times higher than **enclone**, but could have up to 30% higher sensitivity. It is simply not possible to know how much of that 30% comes from true joins, rather than false joins. We do not have an opinion about this.

#### Caveats

1. The error rate is based on interdonor comparisons. As each individual’s exposure to antigens is unique, and each exposure may elicit multiple independent responses, we think that the true error rate for intradonor comparisons (which is what matters) is likely higher.
2. Both the reported error rate and intradonor merges are based on accounting of cell pairs. Thus single true or false joins affect these statistics quadratically. A small number of events can significantly affect the reporting (**Figure 1**).
3. We did not test non-default settings for partis, which might improve performance. Its default setting is deliberately calibrated for high sensitivity at the expense of some specificity.

## Viewing clonotypes

Over the next several sections, we describe some of the visualization capabilities of **enclone**. Considerable help and feature documentation are available through the **enclone** site. As this paper is not intended primarily as a manual, we refer the reader to bit.ly/enclone for specific instructions on use. Help is also available at the command line by typing simply `enclone`.

We think readers will also find **enclone** useful as a tool for thinking about what a clonotype actually is. **enclone** provides a compact view of a clonotype, which we explain in **Figure 2**. Readers who are so inclined can follow along interactively, at least on Mac and Linux computers. To do this, you can type one line to install^5^ **enclone**, following the instructions at bit.ly/enclone, and then type the given command. The command to replicate **Figure 2** is

~~~
enclone BCR=123085 CDR3=CTRDRDLRGATDAFDIW.^6^
~~~

The display employs two types of compression to render it accessible to the eye:

- In *vertical* compression, cells having identical transcripts are combined into what we call **exact subclonotypes**^7^. These appear as single table rows in the display.
- In *horizontal* compression, to represent the full transcript, we show amino acids colored by codon, thus using three-fold less space than bases, and we show only selected amino acids, in a configurable fashion (more detail below).

Notice in the display the line “reference”, which refers to what we call the **universal reference**, the reference sequence provided by the user that approximates the V(D)J sequences present in the species. Conversely the line “donor ref” refers to the **donor reference**, an approximation to the germline sequences possessed by the donor for a particular sample, and which **enclone** computes as a core component of the clonotyping algorithm.

Now we can describe the default configuration for which amino acids are to be shown: it is those lying in the CDR3 region or for which some nucleotide in some cell is different from the universal reference. In the display, amino acids are colored by their originating codon, using a seven-color palette, in such a way that different codons for the same amino acid are assigned different colors. This allows one to see synonymous changes that represent an evolutionary difference, even though they do not affect protein function.

### § Seeing an evolutionary history of a clonotype

Adding the argument `TREE` to an **enclone** command causes a phylogenetic tree (see <u>References</u>) to be displayed, **Figure 3**.

Note the seven “dot” columns that occur outside the CDR3. All happen to be on the heavy chain in this example. Six of them (at amino acid positions 2, 7, 8, 59, 68 and 105) represent codon differences from the donor reference, shared by every cell in the clonotype. (The number of base differences might be larger.) For example, at position 2, there is a synonymous change at a histidine (H). These shared differences tell us that the clonotype evolved substantially, as represented by the first edge in the tree, before giving rise to the cells observed in the sample obtained from the donor. We note that shared mutations as shown above also “power” the clonotyping algorithm, as explained later.

### § Generating output for use with other programs

Sometimes the most effective way to use **enclone** is as a front end for other programs, perhaps written in R or Python (or even Rust). For such purposes, instead of pretty output, intended for direct viewing, one can use **enclone** to generate *parseable* data in CSV format. Doing this requires selection of the desired fields (columns). The relevant **enclone** documentation pages are the commands [enclone help lvars](https://10xgenomics.github.io/enclone/pages/auto/help.lvars.html), [enclone help cvars](https://10xgenomics.github.io/enclone/pages/auto/help.cvars.html), [enclone help parseable](https://10xgenomics.github.io/enclone/pages/auto/help.parseable.html); you can also view the [enclone variable inventory](https://10xgenomics.github.io/enclone/pages/auto/variables.html).

We provide an example of an **enclone** command and its anatomy. Consider the following command:

~~~
enclone BCR=123085 CHAINS_EXACT=2 NOPRINT POUT=put-your-filename-here
PCOLS=group_id,exact_subclonotype_id,cdr1_aa1,cdr2_aa1,cdr3_aa1,cdr1_aa2,
cdr2_aa2,cdr3_aa2
~~~

This command supposes that one wants to analyze the amino acid sequences of all three complementarity-determining regions CDR1, CDR2 and CDR3, for all antibodies in the given BCR dataset for which exactly one heavy and light chain were observed. Details of the command are explained below:

- The argument `CHAINS_EXACT=2` selects antibodies exhibiting exactly two chains (one heavy and one light; the HH and LL cases are filtered out).
- The arguments `NOPRINT` and `POUT=put-your-filename-here` turn off the standard pretty output and send parseable output to the given file.
- The CDR sequences are selected for output using `PCOLS=cdr1_aa1,cdr2_aa1,cdr3_aa1,cdr1_aa2,cdr2_aa2,cdr3_aa2`. The suffixes `1` and `2` are the chain indices in the clonotypes, and because of our simplifying assumption, `1` corresponds to the heavy chain and `2` to the light chain. In TCR data one would get TRB, TRA, in that order.
- Adding the variables `group_id` and `exact_subclonotype_id` to `PCOLS` exports the clonotype and exact subclonotype identifiers to the parseable output.

In the output file `put-your-filename-here`, you will find this output from the full command:

~~~
group_id,exact_subclonotype_id,cdr1_aa1,cdr2_aa1,cdr3_aa1,cdr1_aa2,cdr2_aa2,cdr3_aa2
1.1, GFTFSDY,STNGGN,CVKDRVTGTITELDYW,SGDKLGDKYAC,QDTKRPS,CQAWDSSAGVF
2.1, GFTFGDY,RSKAYGGT,CTRDRDLRGATDAFDIW,RASQSVSSYLA,DASNRAT,CQQRSNWPPSITF
2.2, GFTFGDY,RSKAYGGT,CTRDRDLRGATDAFDIW,RASQSVSSYLA,DASNRAT,CHQRSNWPPSITF
2.3, GFTFGDY,RSKAYGGT,CTRDRDLRGATDAFDIW,RASQSVSSYLA,DASNRAT,CQQRSNWPPSITF
2.4, GFTFGDY,RSKAYGGT,CTRDRDLRGATDAFDIW,RASQSVSSYLA,DASNRAT,CQQRSNWPPSITF
…
~~~

### § Seeing the junction region

**enclone** can display the junction region of immune receptors. For IGH and TRB, it can find an optimal D region, allowing for the possibility of VDDJ (see <u>References</u>). We note that because junction regions undergo extensive changes during recombination, there is not enough information to correctly assign D regions in all cases. Some are wrong. Moreover, in the display of the junction region, it is not possible to know the actual history of insertions and deletions, and thus what we display is just one possible version of this history. **Figure 4** shows an example of VDDJ.

We were curious about the longest heavy chain CDR3 sequences in the data. In order to find these, we use the following command:

~~~
enclone BCR=@test BUILT_IN CHAINS_EXACT=2 CHAINS=2 POUT=stdout
PCOLS=cdr3_len1,d1_name1,datasets,cdr3_aa1,const1,dref NOPRINT | sort −n |
tail −10 | tac
~~~

which (see previous section) uses **enclone** to generate parseable output, then analyzes that in a trivial fashion. This is a large computation. Note also that the same sort of information can be rapidly obtained from the file per_cell_stuff that is [here](https://figshare.com/ndownloader/files/34846929) and is the output of a similar but more comprehensive command using BCR=@test.

We put the resulting information in **Table 2**. Unsurprisingly, these were enriched for use of two D genes; we also observed one clonotype with a fragment of the *GAB3* gene in CDRH3. Many such clonotypes are class-switched, which is unsurprising given that long antibodies are known to be generated and retained during infection. We show the selection bias towards class-switched VDDJ clonotypes as CDRH3 length increases in **Figure 5**.

**Table 2.**
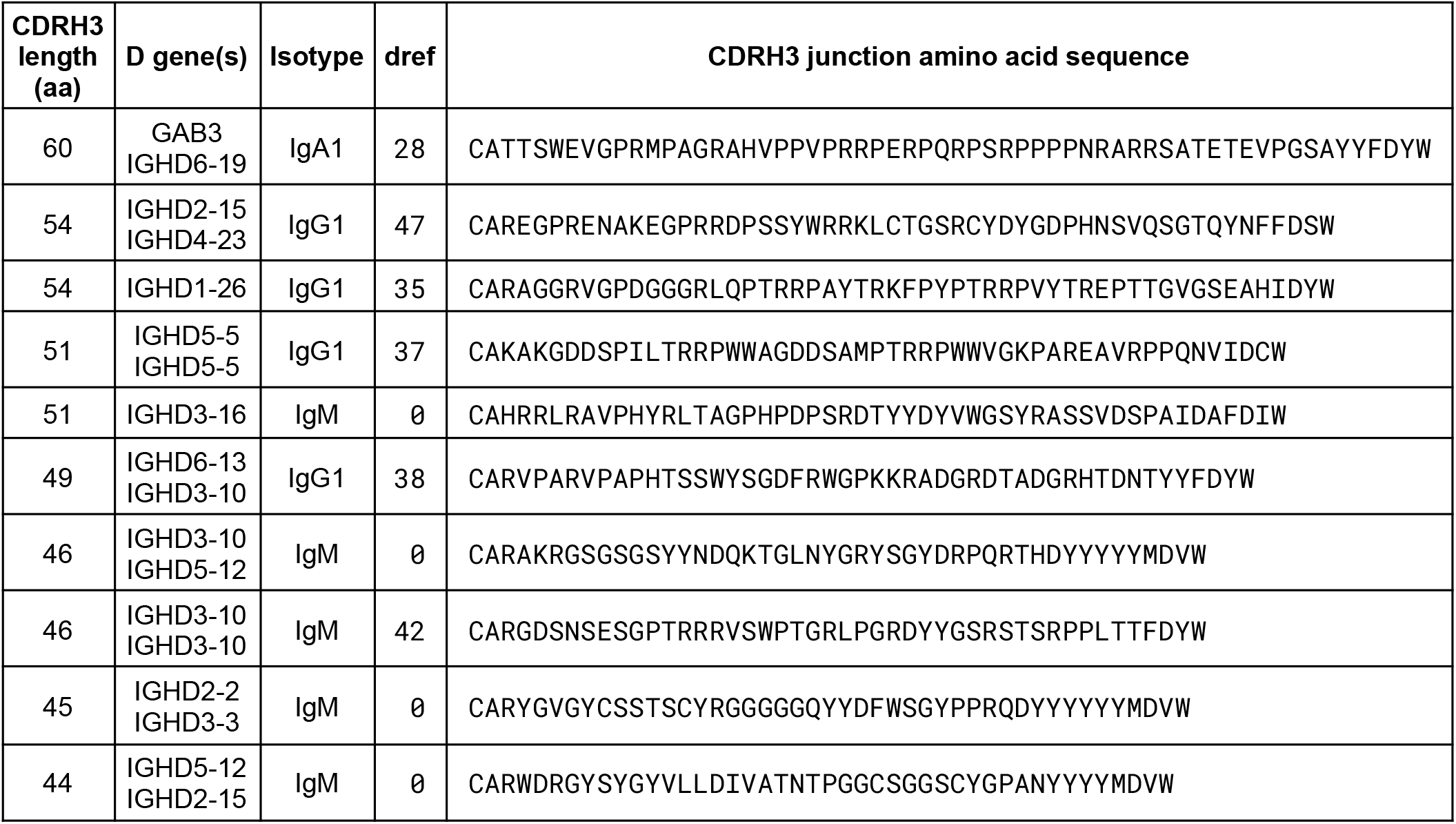
Longest CDRH3 sequences in the test data. We analyzed the lengths of the CDRH3 sequences that occur amongst cells having exactly one heavy and light chain. The table displays the lengths of the 10 longest CDRH3s, along with the D gene(s) that were used. The longest example contains a genomic DNA insert from *GAB3*, specifically 4915:4997 of NG_012834.2. Unsurprisingly, the longest observed junctions are enriched in VDDJ and non-canonical VDJ rearrangements, and class-switched, implying that they have contributed to a memory immune response. The dref column is “donor reference distance”, the inferred number of somatic mutations outside the junction region (totalled over both chains).

### § Detecting insertions and deletions in clonotypes

**enclone** can detect and display SHM insertions and deletions in clonotypes (**Figure 6**).

The algorithm can only find one indel, and only when it occurs *before* the junction region. Although SHM indels can occur inside or after the junction, these appear to be exceptionally rare. We are not aware of a case where such an event has been definitively assigned to SHM, rather than junction rearrangement [Yu 2014], however **Figure 7** may provide one such example.

### § Visualizing all the clonotypes

**enclone** can display a plot showing all the clonotypes in the entirety of clonotypes it computes. There are different ways in which this can be accomplished. For example, one can color the clonotypes by their isotype using a *honeycomb plot* (**Figure 8**).

### § Functional grouping of B cell clonotypes

Biologically we expect (with high probability) that all the cells in a B cell clonotype share both a paratopes and an epitope. Thus they are functionally similar. However, functionally similar cells are not necessarily in the same clonotype, and thus there is utility in **functional groups of clonotypes**.

We give some background on this general problem and refer the reader to the <u>References</u> for more extensive literature. Briefly, for a given antigen, one can typically find examples where an antibody in one individual, and an antibody in another individual, are similar by some criteria. Antibodies grouped in such a fashion are then referred to as ***public antibodies*** or ***public clonotypes*** (noting again the different usage of the word clonotype). Examples from many diseases have been identified.

These similarities typically have a general form involving sharing of V/J genes and some percent identity of nucleotides or amino acids within CDR3, all with respect to either just heavy chains, or also light chains. <u>There is extreme variation between the stringency of conditions that are used to define public clonotypes</u>. We give two examples: (1) identical V/J heavy chain genes and identical CDRH3 length and amino acid sequences [Soto 2019] and (2) same CDRH3 length and 70% amino acid identity along them [Murji 2021].

The limitation of all such approaches, and the reason for extreme variation in conditions is simple: true functional groups are in fact very difficult to characterize, and naive criteria as we describe either grossly “under-connect” functionally similar antibodies, or conversely, connect them to functionally unrelated antibodies. <u>Computational determination of functional groups of antibodies is a fundamentally unsolved problem of the field</u>. In that direction, we note the possibility of deducing structural information from sequence information and using it to group antibodies [Raybould 2021].

**enclone** does not solve this problem, but it provides tooling to experiment with existing naive criteria. In the remainder of this section, we describe this tooling and give examples, including from the test data. More specifically, **enclone** provides functional clonotype grouping based on similarity, by several measures, described at [enclone help display](https://10xgenomics.github.io/enclone/pages/auto/help.display.html), with examples given below. The default behavior of **enclone** is to put each clonotype in its own group. Two general types of nontrivial grouping are provided as options:

1. In *symmetric* grouping, the entirety of clonotypes are partitioned into nonoverlapping groups.
2. In *asymmetric* grouping, each clonotype is treated as the “center” of its own group, which can overlap other groups.

We provide examples of each type. In both cases we use the `CDR3=` argument to restrict the clonotypes that are printed. We begin with the following example for symmetric grouping. Suppose we wanted to place together two clonotypes having the same V and J reference sequences, and for at least one cell in each clonotype, CDRH3 amino acid identity of at least 80%, with transitive extension to partition into non-overlapping groups. Then we would use the arguments `GROUP=vj_refname,cdr3_aa_heavy≥80%`. This is exhibited in **Figure 9**. More examples are given later.

Next, we provide the following example for asymmetric grouping. Suppose that for each clonotype, we wish to show its two closest neighbors, by CDR3 Levenshtein edit distance. Then we would use the arguments

~~~
AGROUP AG_CENTER=from_filters AG_DIST_FORMULA=cdr3_edit_distance
AG_DIST_BOUND=top=2.
~~~

This is exhibited in **Figure 10**.

Now we consider what happens if we use even stricter grouping than described earlier, so as to require perfect identity of CDR3 amino acids.

~~~
enclone GROUP=vj_refname,cdr3_aa_heavy≥100%,cdr3_aa_light≥100% BUILT_IN
MIN_GROUP=2 MIX_DONORS LVARS=donors,n MAX_CHAINS=3 BCR=@test
~~~

Here is a detailed dissection of the command:

- `GROUP=vj_refname,cdr3_aa_heavy≥100%,cdr3_aa_light≥100%` -- performs symmetric grouping by identical V and J gene assignments and CDR3 amino acid sequences
- `MIN_GROUP=2` -- suppresses the display of “trivial” groups containing only one clonotype
- `MIX_DONORS` -- allows clonotypes to contain cells from more than one donor (normally suppressed): groups are always allow to contain cells from more than one donor, because this is expected
- `LVARS=donors, n` -- causes the clonotype tables to display the contributing donors and number of cells for each exact subclonotype
- `MAX_CHAINS=3` -- excludes clonotypes having four or more chains, which have a very high probability of representing cell doublets (and which **enclone** by design does not combine into clonotypes).

This very strict definition yielded 813 groups, comprising in total 10,677 cells (less than 1% of the total). However of these groups, only 85 contain cells from more than one donor. We therefore suspect that most of the 813 groups represent false negatives for clonotyping, *i.e*. cases where computed clonotypes could be correctly merged.

### § Published BCR data

We thought it would be helpful for researchers to have direct access to published single-cell BCR datasets, and therefore endeavored to import published single-cell BCR datasets into **enclone**. In general, not all datasets are consented for public access (*e.g*. in dbGaP), and not all have been deposited into public databases in such a way as to facilitate retrieval. This has been a general challenge for the community, and not just for BCR data.

We did find published data from several studies [Setliff 2019; Ramesh 2020; Woodruff 2020; Sokal 2021; Wang 2021] and are very grateful to the authors for having enabled this. All the data were generated using the 10x Genomics Immune Profiling Platform. If one wanted to run **enclone** on all the data using one command, and just see summary statistics, one could use:

~~~
enclone BUILT_IN ACCEPT_REUSE NOPRINT SUMMARY
META=iReceptor/all.meta,kawasaki/all.meta,LIBRA_seq/all.meta
~~~

After filtering, there are in total 309,535 cells. We note that most of these data predate dual indexing and indeed index hopping occurred, resulting in reuse of barcodes between datasets. **enclone** automatically detects this, provides a list of problems, and exits, unless the argument `ACCEPT_REUSE` is provided, as we have done. Anyone using these data should examine the list of problems, which are not entirely attributable to index hopping. For the same reason, we could not use these data to assess clonotyping.

### § Donor reference analysis

**enclone** clonotyping works by first inferring donor reference sequences for each V gene for which there is enough data. We originally built this into the algorithm because it seemed conceptually essential. Now given the test data, we can assess the actual effect. To do this we turned off the donor reference sequence inference and ran the same analysis as in **Table 1**. When we did this we found that the error rate doubled (from 2.65 x 10^-9^ to 5.29 x 10^-9^), while total intradonor merges decreased by 2.3%, indicating lower sensitivity, and confirming our intuition. We note however that the advantage of donor allele finding diminishes as dataset size decreases.

We were curious what alleles were actually found by the algorithm, and to attempt to gain some understanding of this, we examined the details of the alleles found by the algorithm^8^, in relation to the alleles present in the **enclone** and the IMGT reference^9^. To do this we created a matrix which was labeled on one axis by the merged allele set, and on the other by the variant base positions (**Figure 11**). For a given V gene, we took all the alleles and truncated them on the right as needed to achieve the same length.

We observe in this example a case where the four test set donors contain an allele that is not present in IMGT. Across the four donors, we found 21 alleles that were not present in IMGT, and occurred in at least two of the donors. We note that IMGT includes many alleles which differ by indels, whereas **enclone** as implemented does not see these. Thus a new and comprehensive analysis, based on single cell V(D)J data from many donors, might reveal interesting information about the allele distribution in the population. Alternatively whole diploid genomes not obtained from lymphocytes or lymphoblastoid cell lines would provide a more comprehensive representation, including for genes with lower expression.

## Visualizing clonotypes in a GUI

Here we also describe a GUI (graphical user interface) version of **enclone** called **enclone visual** that has been implemented for Mac OS X and Linux (at present). **It is alpha (α) software.** We built it primarily because it makes it easy to view graphical outputs of **enclone** (such as honeycomb plots as above), in conjunction with clonotype tables. However it also enables other convenient features, as we will describe. To use **enclone visual**, see [here](https://10xgenomics.github.io/enclone/pages/auto/visual.html); briefly, one types enclone VIS i n a Terminal window, which causes the **enclone visual** window to open.

**Figure 12** exhibits an example where the **enclone visual** window is opened, and then the command `enclone BCR=123085 HONEY=out=gui,color=var,u_cell2` i s typed in the window. This displays a honeycomb plot showing all cells, colored by the variable `u_cell2`, which is the UMI count for the second chain (relative to the numbering of columns in a given clonotype).

We designed **enclone visual** to be command-driven. That is, one enters **enclone** commands in the **enclone visual** window. Conversely, with small modifications, these commands can be entered into **enclone** itself. We designed this way for two reasons:

- A user can return to a particular state by simply entering the command, which encodes everything.
- A user can employ a workflow in which you experiment in **enclone visual**, then capture the command you want for subsequent use in an automated script.

In the window you can see many buttons. We labeled these with text, rather than abstract symbols, in an effort to shorten the learning curve. There is also a **Help** button that explains the use of all the buttons. For example, clicking on the **Pic** button yields **Figure 13**.

Several of the buttons are of this type—they expand part of the window so as to more effectively exploit the available real estate on-screen.

### § States and sessions

A series of commands can be entered in **enclone visual**, each defining a *state*. Buttons are provided to navigate through past states: in **Figure 12** (above), blank buttons are visible on the right. These populate with up 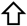 and down 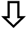 arrows as soon as there are enough states to go forward and backward.

Once one opens the **enclone visual** window, and until one exits, the totality of states created comprise a *session* (akin to a notebook). A session can be saved at any time by pushing the **Save** button near the upper right corner. Pushing the **Archive** button will show the saved sessions, and more, as we describe. Supposing that in **Figure 12**, we pushed the **Archive** button (and had no prior history), we would see this:

<u>Entry 3</u> is the saved session. We will explain the first two entries shortly! First note the ability to create a title and narrative for a given session (instructions at **Expand documentation**), yielding *e.g*.:

The first two entries are special sessions, built into **enclone visual**, called *cookbooks*. At any time, you could open one of those, or open a session that you had previously saved, by checking the **restore** box. You can also curate your list of saved sessions, for example deleting ones that you are no longer interested in.

### § Remote computation (unreleased)

Here we describe a capability built into **enclone**, but which we are withholding, pending feedback, and also experience from the initial release of **enclone visual** (which was approximately concurrent with this paper). Briefly, this capability allows:

1. Use of a remote Linux server as a backend for **enclone visual** computations, and as a common area for dataset storage. The primary advantage of this is that it enables transparent access to shared datasets, without needing to download them.
2. Use of the same server as an intermediary for communications between **enclone visual** users. This allows instantaneous sharing of sessions between users with access to the same server, thus enabling collaboration.

Implementing these features for a given system requires “jumping over” the firewall between the user’s computer and the server (if present). This requires formulation of an appropriate command, and almost certainly requiring new details for particular environments.

### § Seeing the effect of default filtering

Here we use **enclone visual** to exhibit a dramatic example (**Figure 16**) of how **enclone**’s built-in filtering can “clean up” a dataset. On the right, the output of the command `enclone BCR=128040 HONEY=out=gui,color=var,u_cell1 CDR3=CARGGTTTYFISW` is shown. We see **2** cells, colored by their heavy chain UMI counts. On the left, we added the arguments `NUMI NUMI_RATIO` to disable two **enclone** filters, one for filtering by UMI counts and the other for filtering by UMI ratios. Now **122** cells are seen. We hypothesize that only a single *plasma cell* was present, along with fragments of it, which filled droplets (GEMs) and produced an illusion of a much larger population. In other cases (data not shown), we have observed evidence that a single plasma cell disintegrated, leaving only fragments, with no parent in the data.

### § enclone has many optional filters

**enclone** has an extensive range of filters that can be used to select particular clonotypes, documented at `enclone help filter`. For example, we have already seen filters like CDR3=“CARGHYGMDVW|CARDTAVGGDAFDIW”, which select all clonotypes containing one of the CDR3 sequences (and discard the rest). As another example, we can filter by the amount of SHM using the variable `dref`, which is the total number of nucleotide differences between a given contig and the donor reference (outside the junction regions). We can display the effect on isotypes using **enclone visual**, as seen in **Figure 17**.

**Figure 17.**
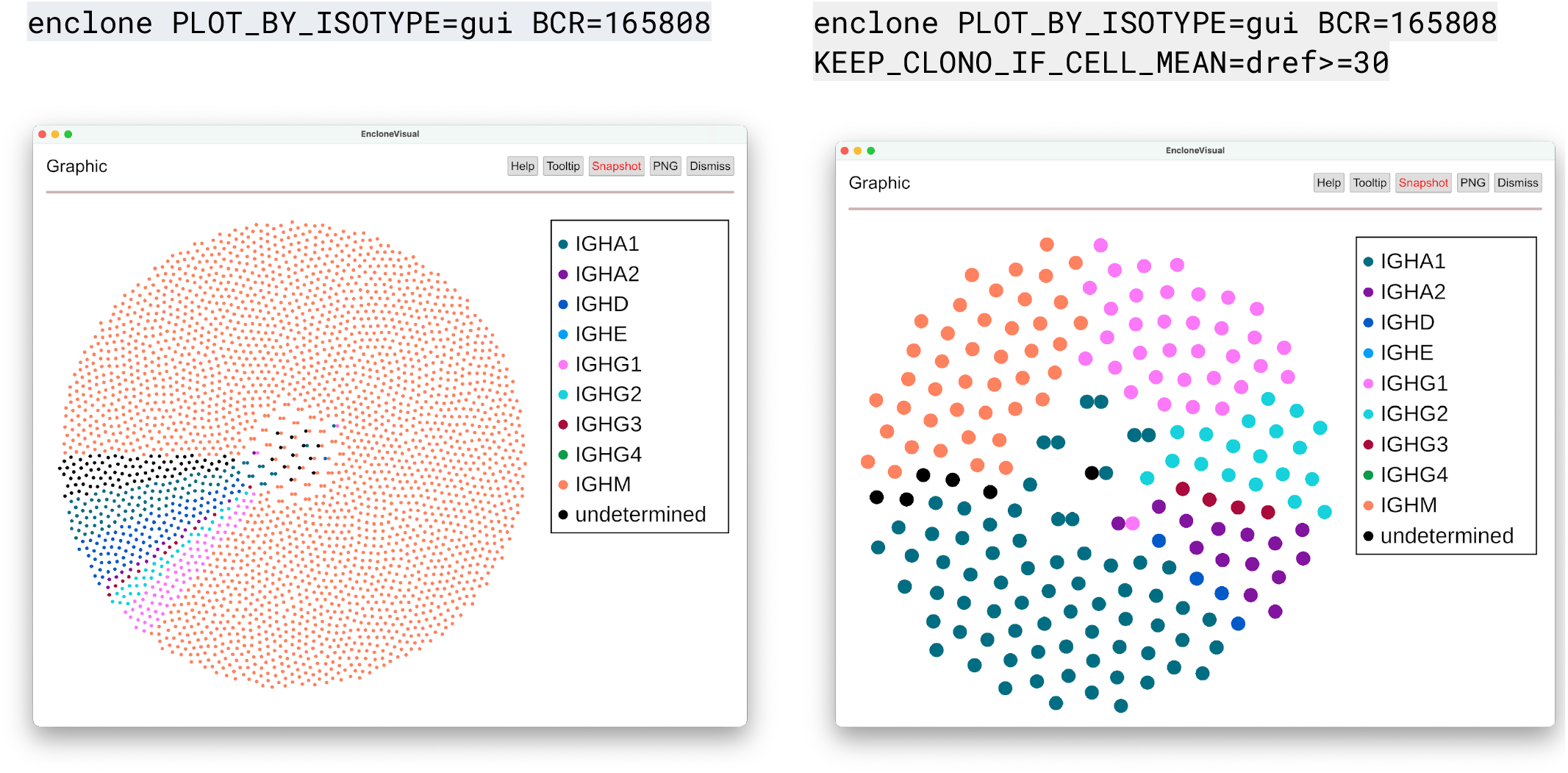
Filtering by SHM shifts isotype distribution. <u>Left panel</u>: honeycomb plot for all cells in a dataset are displayed in **enclone visual**. <u>Right panel</u>: cells are only displayed if they lie in a clonotype for which the mean number of nucleotide differences with the donor reference (outside the junction regions) is at least 30. A far greater fraction of class switching is observed, as expected.

## Methods

### § Reference sequences

Details regarding VDJ reference sequences affect clonotyping and other aspects of biological interpretation. One reason for this is that many V genes are highly similar, so that in some cases V gene assignment is delicate or even arbitrary. Since most clonotyping algorithms use the gene assignments, this can affect results.

As part of this project, we created curated human and mouse V(D)J references that **enclone** can use. Part of this included the following requirements:

- V genes begin with a start codon
- V genes do not have frame shifts
- V genes stop after going exactly one base into the constant region
- Constant regions begin with two bases of a codon.

We optimized **enclone**’s algorithms to work with these reference sequences, and note that performance on other reference sequences can be different. **enclone** can always be forced to use its built-in reference by adding the argument `BUILT_IN`.

As an experiment, we repeated the sensitivity and specificity calculation using the test data (**Table 1**), with default **enclone** settings, but this time used the IMGT reference. When we did this we found that the error rate increased (from 2.65 x 10^-9^ to 3.51 x 10^-9^), and total intradonor merges decreased by 16.5%, suggesting significantly lower sensitivity. These changes might be attributable to defects in the IMGT reference, but also might be attributable to defects in how **enclone** processes the reference.

### § Clonotyping algorithm

#### Exact subclonotypes

**enclone** places cells having essentially identical ..V(D)J.. transcripts in groups called *exact subclonotypes*. We do not require *identical* because 10x technology does not read through the entire C segment, and inconsistently reads the 5’ UTR, so we do not typically know the entire ..V(D)J.. transcript. By *essentially identical* we mean that the V..J sequences are identical, that the reference C segment assignments are the same, and that there is the same distance between the J stop and the C start. This distance is nearly always zero.

Exact subclonotypes that are biologically impossible are discarded: these are those having more than two instances of a given chain type. Therefore in particular, an exact subclonotype can have at most four chains, even though a *clonotype* can have more, see below. Nonproductive transcripts, for example those having frameshift indels, are also discarded.

#### Column structure

In **enclone**, a clonotype is not simply a collection of cells. Rather, it is a *matrix* whose rows are exact subclonotypes and whose columns are chains. This captures the correspondence between chains in different exact subclonotypes. In the canonical case, there are exactly two chains, one heavy and one light (or TRB and TRA), and both chains are present in all cells. In some cells only a single chain is detected. These are included in clonotypes with more chains, so long as the algorithm is confident that the cells belong in the same clonotype. In such cases, the matrix has blank entries for the missing chains (*e.g*. see **Figures 10, 18**), and there are other mechanisms for this, see next.

**Figure 18.**
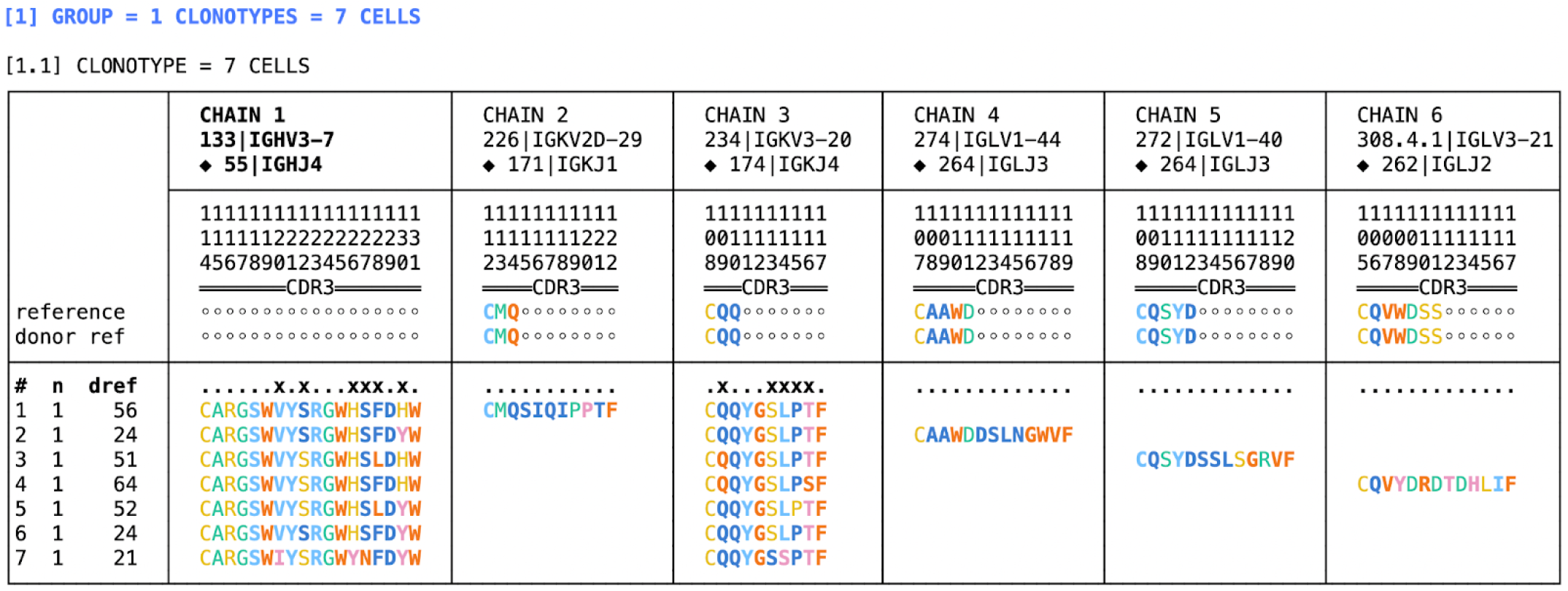
A clonotype having six chains. The start of the output of

~~~
enclone BCR=@test MIN_CHAINS=5 BUILT_IN LVARS=n,dref AMINO=cdr3 FOLD_HEADERS
CVARS=
~~~

is exhibited. There are six chains, even though every exact subclonotype has at most three chains. Cases like this are very rare in *most* datasets. They arise from artifacts in the data or computation rather than actual biology.

#### A clonotype can have many chains

In real data, at some frequency, cells (more precisely GEMs) contain contaminating chains from other cells. This can lead to “false” chains in clonotypes, and this provides a mechanism whereby clonotypes can have more than four chains. Several circumstances could lead to such a state: (1) “too many” cells in a library: (2) excessive background, including from disintegration of plasma cells: (3) concentration of cells in a small number of clonotypes. **enclone** has several filters designed to mitigate these phenomena, which are described later. In the test data, which are “well behaved”, there are 3 clonotypes having 5 chains, 2 having 6 chains (**Figure 18**), and none having more than 6 chains (out of a total of ~1.4M clonotypes).

#### The donor reference

Before clonotyping, **enclone** derives donor reference sequences. Some information about this may be obtained by typing `enclone help how`, and more detail will be provided elsewhere. The same approach was attempted for J segments instead of V segments, but we were unable to improve clonotyping performance with it, and therefore did not include it in the algorithm.

#### Joins to be considered

To build clonotypes, **enclone** considers pairs of exact subclonotypes for joining, *i.e*. inclusion in a common clonotype. Only certain pairs are considered. There are several requirements, one described here and two more below. First, only exact subclonotypes having two or three chains are considered. Clonotypes having one chain are not joined to each other because that would greatly hurt specificity, however there is a separate stage that merges some single-chain clonotypes into clonotypes having more chains. Clonotypes having four chains are not considered because such clonotypes are dominated by cell doublets; joining them would be highly deleterious.

#### Limited indel recognition

Indels could be present in the germline or arise via somatic hypermutations and could theoretically occur at many positions. However, for purposes of clonotyping, **enclone** only detects those that occur before the junction region, and only detects one per transcript. These are limitations of the algorithm. Single indels are detected by alignment to the reference. Insertions are temporarily removed during the clonotyping process, and deletions are replaced by deletion characters (-), so that the lengths of the sequences are restored to their value before the indel occurred.

#### Length identity requirement

For each pair of exact subclonotypes, and for each pair of chains in each of the two exact subclonotypes, for which V..J has the same length for the corresponding chains (after adjustment for a single indel, see above), and the CDR3 segments have the same length for the corresponding chains, **enclone** considers joining the exact subclonotypes into the same clonotype.

#### First test for too many CDR3 mutations

We require that the combined nucleotide identity for CDR3 across the two chains is at least 85%.

#### Divining common ancestry

**enclone** attempts to identify shared somatic hypermutations that would constitute evidence of common ancestry. This is a cornerstone of the algorithm and is the motivation for donor reference computation (above). **enclone** finds shared mutations between exact subclonotypes, that is, for two exact subclonotypes, common mutations from the reference sequence, using the donor reference for the V segments and the universal reference for the J segments. By using the donor reference sequences, most shared germline mutations are excluded, and this is critical for the algorithm’s success.

#### Testing for sufficient shared mutations

We find the probability p that “the shared mutations occur by chance.” More specifically, given d shared mutations, and k total somatic hypermutations (across the two cells), we compute the probability p that a sample with replacement of k items from a set whose size is the total number of bases in the V..J segments, yields at most k – d distinct elements. This probability p is used below in deciding which clonotypes to join.

#### An additional test for too many CDR3 mutations (setup)

Let cd be the number of base positions at which the CDR3 sequences of the two cells differ. We also compute a version cdw of cd that is weighted inversely by CDR3 lengths, as follows. Let cd1 be the number of CDRH3 nucleotide differences, and let cd2 be the number of CDRL3 nucleotide differences. Let n1 be the nucleotide length of CDRH3, and likewise n2 for the light chain. Then cdw = 42 • (cd1/n1 + cd2/n2). Then we set N to 80^cdw^. The higher this number is, the less likely a join is to be correct. See below for how this is used.

#### Key join criteria

Two cells sharing sufficiently many shared differences and sufficiently few CDR3 differences are deemed to be in the same clonotype. That is, the lower p is, and the lower N is, the more likely it is that the shared mutations represent *bona fide* shared ancestry. Accordingly, the smaller pN is, the more likely it is that two cells lie in the same true clonotype. To join two cells into the same clonotype, we require pN ≤ 100,000, or one of the two exceptions described next.

#### First exception to key join criteria

The first exception allows any join that has at least 15 shared mutations *outside* the junction region.

#### Second exception to key join criteria (added after seeing test data)

There is a second exception, which is conceptually based on sharing *inside* the junction region, but whose actual definition is more complicated. It uses only the heavy chain. We first find the apparently optimal D gene, allowing for the case of no D gene, or two D genes. We form the concatenated reference (VDJ, or VJ, or VDDJ). Then we align the junction region on the contig to the concatenated reference. See <u>this section</u>. We score the alignment by counting +1 for each inserted base (present on the contig but not the reference), +1 for each mismatch, and +1 for each deletion, regardless of length. This score is called the heavy chain complexity, denoted hcomp. We then accept any join for which hcomp ≥ 8 and indep ≤ 80, where indep is the number of independent mutations, *i.e*. the number of differences between the two contigs outside the junction region.

#### Mutation imbalance

We reject certain joins that exhibit too great an imbalance in mutations between different regions. Specifically:

- We do not join in cases where there is too high a concentration of changes in the junction region. More specifically, if cd is at least 5 times the number of non-shared mutations outside CDR3 (maxed with 1), the join is rejected.
- If the percent nucleotide identity on heavy chain FWR1 is at least 20 more than the percent nucleotide identity on heavy chain CDR1+CDR2 (combined), then the join is rejected.

#### Other join criteria

- If V gene names are different (after removing trailing *…), and either V gene reference sequences are different, after truncation on right to the same length or 5’ UTR reference sequences are different, after truncation on left to the same length, then the join is rejected.
- We do not join two clonotypes which were assigned different reference sequences, if those reference sequences differ in more than two positions.
- There is an additional restriction imposed when creating two-cell clonotypes: we require that cd ≤ d, where d is the number of shared mutations, as above.
- We do not join in cases where light chain constant regions are different.

### § Filtering algorithms

**enclone** employs a series of filters that are used to remove suspect cells from clonotypes. Without the filters, many clonotypes would be complex tangles with large numbers of chains.

#### Filter outline

We first enumerate the filters (see next page), in the order in which they are applied, and we show their effect on the test data. Importantly, most filters have essentially no effect on the test data. However, every filter has a significant effect on *some* dataset. Datasets vary greatly based on sample preparation and other factors; the filters have been designed to catch a wide range of artifact types.

The filtering steps are intertwined with the clonotyping algorithm. **enclone** provides command line options to selectively turn off these filters, which may be appropriate for certain datasets, or as an exploratory tool to understand why certain cells have been filtered out.

| # | filter name | brief description | percent of test data removed |
| --- | --- | --- | --- |
| 1 | cell filter | remove barcodes not called cells in Cell Ranger VDJ pipeline | (not in accounting) |
| 2 | max contigs filter | remove barcode having more than four contigs | 0.000 |
| 3 | graph filter | remove some exact subclonotypes that appear to be background | 0.066 |
| 4 | cross filter | use cross-library information to remove spurious exact subclonotypes | 0.157 |
| 5 | barcode dup filter | remove duplicate barcodes within an exact subclonotype | 0.000 |
| 6 | whitelist filter | remove rare artifacts arising from gel bead contamination | 0.002 |
| 7 | foursie filter | remove some four-chain clonotypes that might represent doublets | 0.076 |
| 8 | improper filter | remove exact subclonotypes having multiple chains, all of the same type | 0.182 |
| 9 | weak onesie filter | disintegrate some single-chain clonotypes into single cells | 0.000 |
| 10 | UMI filter | remove some B cells having very low UMI counts | 0.780 |
| 11 | UMI ratio filter | remove some B cells having very low UMI counts, relative to clonotype | 0.000 |
| 12 | GEX filter | remove cells called by VDJ pipeline but not by GEX pipeline | 0.000<sup>10</sup> |
| 13 | doublet filter | remove some barcodes that appear to represent doublets | 1.770 |
| 14 | signature filter | remove barcodes that may be contaminants, based on chain signature | 0.035 |
| 15 | onesie merger | prevent merger of some single-chain clonotypes into other clonotypes | 0.000 |
| 16 | weak chain filter | remove cells having a chain that is probably spurious | 0.050 |
| 17 | quality filter | filter out exact subclonotypes having a position with low quality scores | 0.160 |
|  |  | **TOTAL** | **3.278** |

##### 1. Cell filter

This filter removes any barcode that was not declared a cell by the upstream (Cell Ranger) VDJ pipeline.

##### 2. Max contigs filter

Remove any barcode having more than four productive contigs. The cell filter also removes these contigs, so this filter only has an effect if the cell filter is turned off.

##### 3. Graph filter

This filter removes exact subclonotypes that by virtue of their relationship to other exact subclonotypes, appear to arise from background mRNA or a phenotypically similar phenomenon.

##### 4. Cross filter

If you specify that two or more libraries arose from the same origin (i.e. cells from the same tube or tissue), then by default enclone will "cross filter" so as to remove expanded exact subclonotypes that are present in one library but not another, in a fashion that would be highly improbable, assuming random draws of cells from the tube. These are believed to arise when a plasmablast or plasma cell breaks up during or after pipetting from the tube, and the resulting fragments seed GEMs, yielding expanded ‘fake’ clonotypes containing mRNA from real single plasma cells.

If a V..J segment appears in exactly one dataset, with frequency n, let x be the total number of productive pairs for that dataset, and let y be the total number of productive pairs for all datasets from the same origin. If (x/y)^n^ ≤ 10^-6^, *i.e*. the probability that assuming even distribution, all instances of that V..J ended up in that one dataset, delete all the productive pairs for that V..J segment that do not have at least 100 supporting UMIs.

##### 5. Barcode duplication filter

If the same barcode appears more than once within a given exact subclonotype, then all instances of the barcode within that exact subclonotype are removed.

##### 6. Whitelist filter

This filters out certain rare artifacts from contamination of oligos on gel beads.

##### 7. Foursie filter

Foursie exact subclonotypes (i.e. those having four chains) are highly enriched for cell doublets. Deleting them all might be justified, but because it is hypothetically possible that sometimes they represent the actual biology of single cells, we do not do this. However we never merge them with other exact subclonotypes, and sometimes we delete them, if we have other evidence that they are doublets.

For each foursie exact subclonotype, **snclone** looks at each pair of two chains within it (with one heavy and one light, or TRB/TRA), and if the V..J sequences for those appear in a twosie exact subclonotype (i.e. one having two chains) having at least ten cells, then the foursie exact subclonotype is deleted, no matter how many cells it has.

##### 8. Improper filter

Filter out exact subclonotypes having more than one chain, but all of the same type. For example, the filter removes all exact subclonotypes having two TRA chains and no other chains.

##### 9. Weak onesie filter

Disintegrate certain untrusted clonotypes into single cell clonotypes. The untrusted clonotypes are onesies (single-chain clonotypes) that are light chain or TRA and whose number of cells is less than 0.1% of the total number of cells. This operation reduces the likelihood of creating clonotypes containing cells that arose from different recombination events.

##### 10. UMI filter

**enclone** filters out B cells having low UMI counts, relative to a baseline that is determined for each dataset.

The motivation for this filter is to mitigate illusory clonotype expansions arising from fragmentation of plasma cells or other physical processes (not all fully understood). These processes all result in “cells” having low UMI counts, many of which do not correspond to intact real cells. Illusory clonotype expansions are generally infrequent, but occasionally cluster in individual datasets.

Nomenclature: for any cell, find the maximum UMI count for its heavy chains, if any, and the maximum for its light chains, if any. The sum of these two maxima is denoted umitot.

The algorithm for this filter first establishes a baseline for the expected value of umitot, for each dataset taken individually. To do this, all clonotypes having exactly one cell and exactly one heavy and light chain each are examined. If there are less than 20 such cells, the filter is not applied to cells in that dataset. Otherwise, let n_50% denote the median of the umitot values for the dataset, and let n_10% the 10th percentile. Let

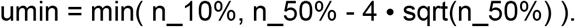

This is the baseline low value for umitot. The reason for having the second part of the min is to prevent filtering in cases where UMI counts are sufficiently low that Poisson variability could cause a real cell to appear fake.

Next we scan each clonotype having at least two cells, and delete every cell having umi_tot_ < u_min_, with the following qualifications:

- Let k be the number of cells to be deleted in a clonotype having n cells. Then we require that for a binomial distribution having p = 0.1, the probability of observing k or more events in a sample of size n is less than 0.01. The more cells are flagged in a clonotype, the more likely this test is satisfied, which is the point of the test.
- If every cell in a clonotype would be deleted, then we find its exact subclonotype having the highest sum for umi_tot_, summing across its cells. Then we protect from deletion the cell in this exact subclonotype having the highest umi_tot_ value. We do this because in general even if a clonotype expansion did not occur, there was probably at least a single *bona fide* cell that gave rise to it.

A better test could probably be devised that started from the expected distribution of UMI counts. The test would trigger based on the number and improbability of low UMI counts. The current test only considers the number of counts that fall below a threshold, and not their particular values.

##### 11. UMI ratio filter

**enclone** filters out B cells having low UMI counts, relative to other UMI counts in a given clonotype.

Each clonotype is examined. We mark a cell for possible deletion if the VDJ UMI count for some chain of some other cell in the clonotype is at least 500 times greater than the total VDJ UMI count for the given cell. Let n be the number of cells in the clonotype. Let k be the number of cells marked for deletion. Then we delete the marked cells if for a binomial distribution having p = 0.1, the probability of observing k or more events in a sample of size n is less than 0.01.

##### 12. GEX filter

If gene expression and/or feature barcode data are provided to **enclone**, and if a barcode is called a cell by the VDJ part of the upstream (Cell Ranger) pipeline, but not called a cell by the gene expression and/or feature barcode part, then **enclone** ignores the barcode.

##### 13. Doublet filter

This removes some exact subclonotypes that appear to represent doublets (or possibly higher-order multiplets). Two exact subclonotypes in a given clonotype have the same *chain signature* if they have the same column entries in the clonotype. For example, in **Figure 18**, five chain signatures are present: one for each of exact subclonotypes 1-4, and one for the three exact subclonotypes 5-7.

The algorithm first computes *pure* subclonotypes. This is done by taking each clonotype and breaking it apart according to its chain signature. All the exact subclonotypes having the same chain signature define a pure subclonotype.

In the simplest case, where the clonotype has two chains, the clonotype could give rise to three pure subclonotypes: one for the exact subclonotypes that have both chains, and one each for the subclonotypes that have only one chain.

The algorithm then finds triples (p0, p1, p2) of pure subclonotypes, for which the following three conditions are all satisfied:

- p0 and p1 share an identical CDR3 DNA sequence
- p0 and p2 share an identical CDR3 DNA sequence
- p1 and p2 do not share an identical CDR3 DNA sequence.

Finally, if 5 • ncells(p0) ≤ min(ncells(p1), ncells(p2)), the entire pure subclonotype p0 is deleted. After all these operations are completed, the algorithm finds cases where a clonotype can be broken into two or more smaller clonotypes, with no shared chains between them, and breaks up the clonotype in such cases.

##### 14. Signature filter

This filter removes some exact subclonotypes that appear to represent contaminants, based on their chain signature. This filter sometimes breaks up complex clonotypes having many chains and representing multiple true clonotypes that are glued together into a single clonotype via exact subclonotypes whose constituent barcodes do not arise fully from single cells.

The algorithm uses some terminology described at doublet filtering, above. Given a pure subclonotype p having at least two chains, if the total cells in the two-chain pure subclonotypes that are different from it but share a chain with it is at least 20 times greater than the number of cells in p, then p is deleted.

##### 15. Onesie merger

**enclone** merges certain onesie clonotypes into clonotypes having two or more chains. The onesie merger filter precludes these merges if the number of cells in the onesie is less than 0.01% of the total number of cells.

##### 16. Weak chain filter

If a clonotype has three or more chains, and amongst those there is a chain that appears in a relatively small number of cells, we delete all the cells that support that chain. The precise condition is that the number of cells supporting the chain is at most 20, and 8 times that number of cells is less than the total number of cells in the clonotype.

##### 17. Quality filter

**enclone** filters out exact subclonotypes having a base in V..J that looks like it might be wrong, based on quality scores.

**enclone** finds bases which are not Q60 for a barcode, not Q40 for two barcodes, are not supported by other exact subclonotypes, are variant within the clonotype, and which disagree with the donor reference. An exact subclonotype having such a base is deleted.

## Supplementary information

### § Authorship confirmation and material sharing

All authors have read, written, and approved the manuscript. This manuscript has not been accepted for publication or published elsewhere. This manuscript is made available on bioRxiv under the CC-BY-ND license; bioRxiv is granted a perpetual, non-exclusive license to display this manuscript.

## § Acknowledgements

This work was fundamentally enabled by the invention of a powerful technology for single cell V(D)J sequencing by 10x Genomics, and by the generation over a period of years of an extraordinarily large training set as a by-product of refining that technology, and by the generation of a large test set just for this work. We thank the entire company for this.

We thank our customers Albert Vilella and Ganesh Phad for many insightful questions that helped shape **enclone**. We thank Pat Marks for proposing V(D)J work years ago, that led ultimately to this project, and for many interesting ideas along the way. We thank Mike Stubbington for user feedback and being a guinea pig, a series of stimulating conversations, and for arranging data generation critical to this work. We thank Sreenath Krishnan for lending his deep expertise on numerous occasions and for his code contributions.

We thank Shaun Jackman for the innumerable occasions on which he volunteered his help and rescued us from an otherwise unsolvable problem. We thank Adam Azarchs and Luiz Irber for lending their extensive software expertise. We thank Nima Mousavi for adding gamma-delta TCR support.

We thank the 10x Genomics support team including Rachana Jain, Nisha Pillai, and Nur-Taz Rahman for asking so many questions and bringing us use cases we had not thought of. We thank Jessica Hamel and Alvin Liang for enduring a series of painful sessions in which they got us started on visualization, and for code they wrote; also Lance Helper, Abou Diop, and Nigel Delaney for their contributions to this. We thank Héctor Ramón for pioneering the Rust crate `iced` that enabled **enclone visual** and Brock (13r0ck) for helping us use it. We thank Keri Dockter for the lovely banner that graces the top of the pages on bit.ly/enclone. We thank Meryl Lewis and Rudy Rico for extensive help with testing machinery.

We also thank members of the Kleinstein laboratory at Yale University, specifically Ken Hoehn, Hailong Meng, and Steve Kleinstein, for their gracious and timely assistance in running the Immcantation Docker container and the associated pipeline correctly. We also wish to thank Nima Nouri, who wrote and maintained SCOPer during his time as a postdoctoral researcher in the Kleinstein laboratory. Similarly, we thank Duncan Ralph and Erick Matsen for their assistance in operating partis correctly, for providing outstanding and unreasonably fast support, and for developing and maintaining partis.

## § Software availability

**enclone** and its source code are available at https://github.com/10XGenomics/enclone. The software is licensed for use in connection with data from 10x Genomics. **enclone** is written entirely in Rust, and uses several Rust crates, created by the developers, that have no restrictions on their use. The same vehicle might be used by new developers who wish to contribute open source code.

## § Reproduction of results

The instructions at bit.ly/enclone may be used to download the *latest* version of **enclone**. At the time you are reading this, it may have been changed, and therefore to reproduce the results of this work verbatim you should instead download the version that was used for this paper, using one of these two links (depending on whether you are using a Linux or Mac computer):

- https://github.com/10XGenomics/enclone/releases/download/v0.5.175/enclone_linux
- https://github.com/10XGenomics/enclone/releases/download/v0.5.175/enclone_macos

depending on whether you are using a Linux or Mac computer. This version corresponds to git hash `c86bd5` of the source code. You would then rename `enclone_linux` or `enclone_macos` to `enclone`.

To generate the first row of **Table 1** (**enclone** run with default parameters), one would run this:

~~~
enclone BCR=@test MIX_DONORS MIN_CHAINS_EXACT=2 NOPRINT BUILT_IN SUMMARY
~~~

On a server having 24+ cores, and depending on its exact configuration, this takes 20-35 minutes. The reported peak memory varied from 120 to 150 GB.

The argument `MIN_CHAINS_EXACT=2` excludes cells for which only one chain was observed. We did this because the merger of such cells into clonotypes by **enclone** was unique amongst the algorithms tested, and therefore turning off this feature facilitated direct comparison.

For MLS: 10 we added the argument `MAX_LOG_SCORE=10`, and similarly for MLS: 15 and MLS: 20.

Testing of the other clonotyping heuristics was more complicated because all the algorithms needed to run on the same cells, and **enclone** filters out cells that appear to be artifacts. Therefore to maintain a completely level playing field, we first ran the alternative algorithm on all the cells, then post filtered by removing cells that were not part of the clonotypes generated by the default **enclone** command. Thus we first ran a command which generated the list of pairs (dataset id, barcode) corresponding to the cells that survived the **enclone** filtering:

~~~
enclone BCR=@test MIX_DONORS MIN_CHAINS_EXACT=2 NOPRINT BUILT_IN
POUT=stdout PCELL PCOLS=datasets_cell,barcode PCOLS_SHOW=dataset,barcode
> post_filter.csv
~~~

For BASIC_H: 85, we then ran this command:

~~~
enclone BCR=@test MIX_DONORS MIN_CHAINS_EXACT=2 NOPRINT BUILT_IN
NALL_CELL POST_FILTER=post_filter.csv JOIN_BASIC_H=85
~~~

where the argument `NALL_CELL` turns off the enclone filtering steps. The other BASIC and BASIC_H algorithms were handled in the same fashion, using instead `JOIN_BASIC=…` for BASIC.

To run and assess SCOPer and partis, we ran this sequence of operations, which use several commands that require compilation of the `enclone` codebase:

1. build_clonotyping_inputs @test, where @test refers to a list of datasets that are available as part of enclone. Users can also specify their own datasets.
2. We ran SCOPer using version 4.3.0 of the Immcantation Docker container. Code to reproduce these analyses is provided at [the enclone GitHub repository](https://github.com/10XGenomics/enclone/tree/master/enclone_paper) using the `autopilot_master.sh` and `autopilot.R` scripts. We ran the SCOPer experiments on an m5a.24xlarge AWS instance with two AMD EPYC 7571 CPUs and a total of 48 cores (96 threads) and approximately 371 GB of RAM. Of note, this is a different machine than we used to benchmark enclone, which was an r5.24xlarge AWS instance with two Intel Xeon Platinum 817M CPUs and a total of 48 cores (96 threads) and 748 GB of RAM. We ran partis using digest 973e400c79a3 of the Docker container on a separate machine. All code used to run both partis and SCOPer as described here is [provided on GitHub](https://github.com/10XGenomics/enclone/tree/master/enclone_paper/src). Both SCOPer and partis have AIRR output modes which generate an AIRR-compatible TSV file that can be assessed using the `assess_clonotyping` binary.
3. enclone BCR=@test MIX_DONORS MIN_CHAINS_EXACT=2 NOPRINT SUMMARY BUILT_IN, which reports the number of inter-donor comparisons for **enclone** on a 1.6M cell publicly available dataset.
4. `assess_clonotyping clones.tsv post_filter.csv cross` (where `cross` is used to calculate the number of inter-donor comparisons)

## § Revision history

1. Apr 21, 2022. This is the first version.

## Footnotes

1 Pronounced ‘en-, klōn.

2 An early reference to this notion of clonotype is [Klinman 1975], “the maintenance of the stimulated B cell’s clonotype (unique individual antibody specificity) in both the antibody forming cell progeny and in subsequent generations of secondary precursor cells”. We note that the term “clonotype” has been used inconsistently (see <u>References</u>). For this work, we always use the definition, or approximations thereof, of “clonotype” we have given.

3 This is the error rate which represents false assignment of cells to clonotypes. Other error types include representation of non-cells as cells, inclusion of contaminating chains in cells, and base errors in chain sequences. We do not attempt to quantify these error types, though minimizing them comprises a major component of the enclone design

4 enclone takes as input a subset of the output files generated by the 10x Genomics Cell Ranger pipeline, as documented here: enclone help input_tech.

5 Use the `medium` option to enable replication of all examples from this paper.

6 Verbatim replicability applies if the **enclone** code has not evolved. See <u>Reproduction of results</u>.

7 An **exact subclonotype** consists of all cells in a clonotype having identical immune receptor transcripts “in principle”; we test for the same number of chains, identical V..J sequences, the same constant region reference sequence, and the same gap between J and C (normally zero); different transcripts emitted by the same cell (notably as in IGHM/IGHD) are not captured by this notion.

8 Adding the argument `DONOR_REF_FILE=filename` causes enclone to dump out the donor reference alleles.

9 Sequences downloaded Jan. 31, 2022.

10 Not applicable because we did not supply gene expression data when we ran **encione** here.

